# RNAi-mediated silencing of SOD1 profoundly extends survival and functional outcomes in ALS mice

**DOI:** 10.1101/2024.06.20.599943

**Authors:** Alexandra Weiss, James W. Gilbert, Iris Valeria Rivera Flores, Jillian Belgrad, Chantal Ferguson, Elif O. Dogan, Nicholas Wightman, Kit Mocarski, Dimas Echeverria, Ashley Summers, Brianna Bramato, Nicholas McHugh, Raymond Furgal, Nozomi Yamada, David Cooper, Kathryn Monopoli, Bruno M.D.C. Godinho, Matthew R. Hassler, Ken Yamada, Paul Greer, Nils Henninger, Robert H. Brown, Anastasia Khvorova

## Abstract

Amyotrophic lateral sclerosis (ALS) is a fatal neurodegenerative condition, with 20% of familial and 2-3% of sporadic cases linked to mutations in the cytosolic superoxide dismutase (SOD1) gene. Mutant SOD1 protein is toxic to motor neurons, making SOD1 gene lowering a promising approach, supported by preclinical data and the 2023 FDA approval of the GapmeR ASO targeting SOD1, tofersen. Despite the approval of an ASO and the optimism it brings to the field, the pharmacodynamics and pharmacokinetics of therapeutic SOD1 modulation can be improved. Here, we developed a chemically stabilized divalent siRNA scaffold (di-siRNA) that effectively suppresses SOD1 expression *in vitro* and *in vivo*. With optimized chemical modification, it achieves remarkable CNS tissue permeation and SOD1 silencing *in vivo*. Administered intraventricularly, di-siRNA^SOD1^ extended survival in SOD1-G93A ALS mice, surpassing survival previously seen in these mice by ASO modalities, slowed disease progression, and prevented ALS neuropathology. These properties offer an improved therapeutic strategy for SOD1-mediated ALS and may extend to other dominantly inherited neurological disorders.

**One sentence summary:** Silencing SOD1 with chemically optimized divalent siRNA profoundly extends the lifespan of G93A mice and prevents neurodegenerative biochemical and behavioral phenotypes.

## INTRODUCTION

Amyotrophic lateral sclerosis (ALS) is a fatal neurogenerative condition characterized by focal and then disseminated weakness, loss of motor function and median age of onset of 50-60 years (*1*). ALS patients typically survive 2-5 years after diagnosis, with respiratory muscle failure being the most common causes of death (*1, 2*). Up to 90% of ALS cases are sporadic, but onset and progression are associated with hundreds of genetic mutations (*3, 4*). The remaining cases are familial, typically transmitted as dominant traits; these arise from mutations in several genes, including *C9orf72* (∼40% of familial cases) (*5, 6*) superoxide dismutase (*SOD1*, 20%) (*7*), followed by fused-in-sarcoma (*FUS*, 5%) (*8, 9*) and transactive DNA binding protein (*TDP43*, 5%) (*10*). Mutations in *SOD1* and *C9orf72* respectively account for ∼1% and 10% of sporadic ALS (*11*).

SOD1 is an antioxidant enzyme that catalyzes the conversion of reactive oxygen species to oxygen and water. The pathophysiology of mutant SOD1 is complex. Many mutant SOD1 proteins show conformational instability and can be induced to aggregate, provoking multiple types of cellular dysfunction (*12*), many mediated by adverse oxidative activity (*13–15*). In transgenic ALS mouse models, knockdown of SOD1 delayed disease onset and preserved neuromuscular junction (NMJ) integrity (*16, 17*). Thus, reducing SOD1 expression in the context of ALS is a promising treatment approach.

In 2023, tofersen, a chemically stabilized antisense oligonucleotide (ASO) that reduces SOD1 mRNA and protein expression, became the first and only FDA-approved drug specifically for SOD1 familial ALS (*18*). Tofersen is pan-SOD1 silencing, targeting both non-mutant and mutant SOD1 alleles. Although the clinical trial failed to reach its clinical endpoint (improvement in ALSFRS-R score) (*19, 20*), tofersen was ultimately approved after results during the open-label extension showed that disease progression was stabilized in patients who received earlier initiation of tofersen treatment versus those who received delayed treatment (*21*). Tofersen also reduced levels of the neuronal injury marker, neurofilament light chain (Nfl) (*22*), suggesting attenuation of neuronal loss following SOD1 reduction. While tofersen represents a profound advance in therapy for SOD ALS, tofersen’s clinical success is potentially limited by unfavorable pharmacokinetic and pharmacodynamic (PK/PD) properties of its GapmeR ASO (modified RNA flanks with an unmodified DNA core) design including limited on-target SOD1 lowering, frequent administration, and a high dose regimen. A more robust treatment that can halt, prevent, or reverse SOD1-mediated familial ALS would be preferable for patients with this devastating condition.

RNA interference (RNAi)-based therapeutics, including small interfering RNA (siRNA), offer a robust strategy to potently, specifically, and safely silence therapeutic gene targets.

Whereas the siRNA sequence determines gene target specificity, the chemical scaffold (i.e., modification pattern and structure) of the siRNA defines stability and delivery *in vivo*. In fact, SOD1 lowering by siRNA conjugated to an ASO was recently demonstrated to extend survival of ALS mice *in vivo* demonstrating that siRNA-ASO conjugated compounds drive more potent and longer-lasting silencing of SOD1, translating into measurable disease-modifying effects (*23*).

The 2’-O-hexadecyl (C16) lipid conjugate (*24*) and the divalent siRNA scaffold (*25*) are currently the two most advanced siRNA scaffolds for central nervous system (CNS) delivery. In the divalent scaffold, two siRNA are linked through their passenger strand (*25, 26*). The increase in molecular size of divalent siRNA (di-siRNA) compared to mono-valent or smaller entities reduces the rate of cerebrospinal fluid (CSF) clearance to facilitate higher CNS retention and broader distribution, potentially enhancing the efficiency of neuronal uptake (*27*). When fully chemically modified to increase nuclease resistance, enhance biodistribution, and prevent immunoreactivity, di-siRNA achieves at least 6 months of target gene silencing in rodent and non-human primate (NHP) CNS following a single injection (*25*). Fully chemically modified siRNAs are widely used in clinical practice (*28–30*), with more than five chemically modified siRNA drugs being FDA-approved for liver-related indications (*28*).

Here, we identified and characterized a potent, chemically modified di-siRNA targeting SOD1, called di-siRNA^SOD1_123^. A single injection chemically-optimized di-siRNA^SOD1_123^ into the CSF of the mutant SOD1 overexpression mouse model G93A, significantly extended lifespan past 250 days, outperforming tofersen. Repeat di-siRNA^SOD1_123^ injections (every 8 weeks) further extended G93A lifespan past 300 days. Compared to no treatment or control treatment, di-siRNA^SOD1_123^ prevented neuronal loss in the motor cortex and lumbar spinal cord, maintained cortical and lumbar synapse integrity, and prevented NMJ loss. The optimized di-siRNA^SOD1_123^ characterized in this study presents a new modality for treatment of SOD1 ALS.

## RESULTS

### Extensive screening identifies a potent chemically stabilized SOD1-targeting siRNA

To identify active siRNAs that silence human SOD1, we designed and screened a panel of siRNA sequences that target different regions of SOD1 mRNA (e.g., open reading frame, 5’ and 3’ untranslated region) using a previously validated bioinformatics algorithm to rank mRNA target regions (*31*). Every sequence was fully chemically modified (Figure 1A, B) to stabilize the nucleic acid backbone against nucleases which is essential for siRNA activity *in vivo.* Chemical modifications of siRNA include stabilization of the nucleic acid backbone using phosphorothioate (PS), and sugar modifications to the 2’ position, most commonly 2’ methyl (2’OMe) and 2’ fluoro (2’F). The compounds were asymmetric and conjugated to cholesterol, which supports internalization without transfection reagents mimicking uptake *in cellula* through endocytosis (*32*). The exact sequence and chemical modification pattern used is shown in the Table S1. siRNAs were named based on their targeting position in the SOD1 mRNA.

**Fig. 1.**
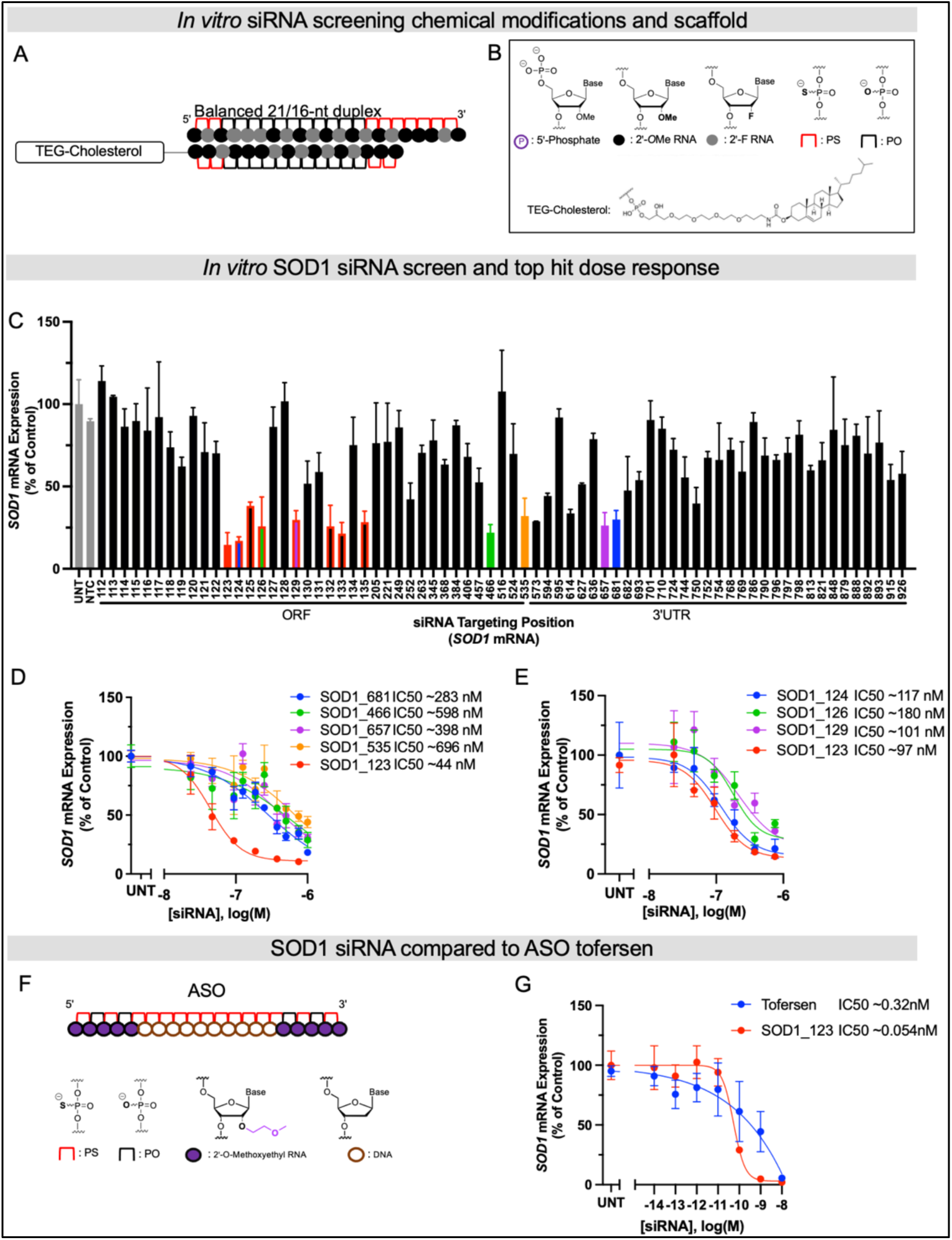
Systematic screening of chemically modified siRNAs targeting SOD1 mRNA identifies a highly functional compound. (**A** and **B**) Schematics of chemical modifications and siRNA scaffolds tested in screen. **(C)** In vitro screen of SOD1 siRNAs in HeLa cells. HeLa cells were treated (by passive uptake) with a panel of siRNAs (1.5 µM) for 72 hours. **(D)** 7-point dose-response curves of lead siRNA compounds identified in primary screen in C for 72 hours. **(E)** 7-point dose-response curves of lead compounds from siRNA “walk” screen marked in red in the bar graph in C. **(F)** Schematic of ASO Tofersen and its modifications. **(G)** 7-point dose-response curves of Tofersen and SOD1_123 with lipid-mediated uptake. All mRNA levels were measured using QuantiGene (n=3, bar is average n=3 ± SD) IC50 values were calculated using the nonlinear least squares method (GraphPad Prism). NTC (nontargeting control siRNA), UNT (untreated).

Primary screening in human HeLa cells identified multiple active siRNA “hits”, with effective SOD1 mRNA silencing up to ∼80% (Figure 1C, colored bars). The IC50 values were determined for the top five hits (SOD1_123, 466, 535, 657, and 681) using 7-point dose-response curves. SOD1_123 was the most potent compound, with an IC50 of 44 nM (Figure 1D). To explore whether SOD1_123 is in a region highly accessible to RNAi machinery, we performed a “walk” around the SOD1_123 target site to generate a new panel of siRNAs (spanning positions 123-133). This walk panel was screened in HeLa cells, with seven additional active sequences being identified (Figure 1C, bars outlined in red). However, SOD1_123 remained the most potent compound in a 7-point dose response assay (Figure 1E).

To benchmark the potency of the lead siRNA, SOD_123, against tofersen (Figure 1F), we synthesized tofersen as described previously (*21, 33*) and performed a 7-point dose response assay in HeLa cells. Because ASOs do not efficiently self-internalize *in vitro* at the range of concentrations and timepoint (72 hours) used in this assay, we used lipofection to deliver both tofersen and SOD_123. SOD_123 was six times more potent than tofersen (IC50 of 54 pM vs. 320 pM, respectively, Figure 1G).

An essential step toward clinical development of siRNA therapeutics is to evaluate their safety and efficacy in animal models with tissue characteristics resembling those in humans, such as NHPs. Identifying human SOD1-targeting siRNA with cross-species activity for testing in NHPs would accelerate preclinical development of siRNAs for SOD1 ALS. To determine the cross-reactivity of our lead siRNA, SOD_123, we evaluated potency in the NHP cell line, LLC-MK2 (Figure S1). The guide sequence of SOD1_123 is fully complementary to human SOD1 mRNA but has a mismatch at position 13 against NHP SOD1 mRNA (Figure S1). A single mismatch can abolish siRNA activity or have no detectable impact (*34*). We therefore converted the mismatched nucleotide in SOD1_123 to be fully complementary to the NHP SOD1 targeting site, creating a new siRNA, SOD1_NHP. We delivered SOD1_123 or SOD1_NHP to LLC-MK2 cells and human cells (Figure S1) and performed 7-point dose-response assays. Both SOD1_123 and SOD1_NHP exhibited good potency in NHP cells, suggesting the single mismatch is well tolerated. Thus, SOD1_123 can be used for preclinical studies in NHP.

### di-siRNA^SOD_123^ extends survival and slows disease progression in an ALS mouse model

We next sought to use our efficacious, potent, and fully chemically stabilized lead, SOD1_123, to understand the impact of SOD1 silencing on disease progression. To achieve CNS delivery, we synthesized SOD1_123 in a di-siRNA scaffold using the same fully chemically modified designs used in screening (di-siRNA^SOD1_123^) (Figure 2A-C).

**Fig 2.**
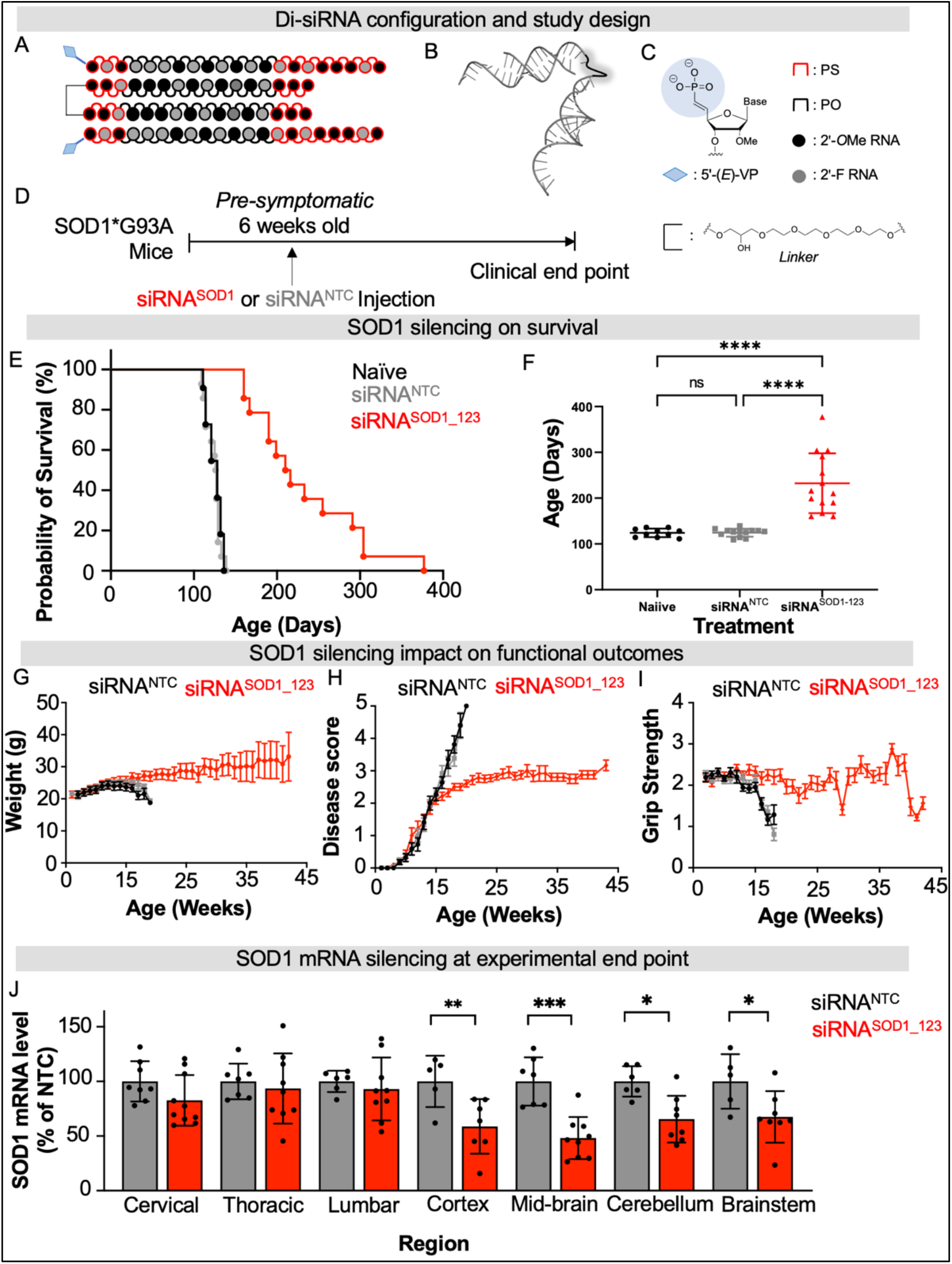
Di-siRNA^SOD_123^ extends lifespan of SOD1 G93A ALS mice. (**A-C**) Di-siRNA^SOD_123^ scaffold (**A**), PyMOL structure (**B**) and chemical modifications used (**C**). (**D**) Study design. (**E**) Kaplan-Meier survival curve of naïve (black), non-targeting control (NTC, gray) treated, or di-siRNA^SOD1_123^ treated (red) G93A mice. (**F**) Plotted age at death for naive (black), NTC (gray) or di-siRNA^SOD1_123^ (red) treated mice. (**G-I**) Naïve (black), non-targeting control (NTC, gray) treated, or di-siRNA^SOD1_123^ treated (red) G93A mice weight (**G**), disease score (**H**), or grip strength (**I**) monitored weekly until clinical endpoint. (**J**) SOD1 mRNA in NTC (gray) or di-siRNA^SOD1_123^ (red) treated mice. Tissue collection occurred at experimental endpoint. * p < 0.05, ** p< 0.01, *** p < 0.001, **** p < 0.0001, Two-way ANOVA with multiple comparisons.

To evaluate the impact of *in vivo* RNAi-mediated SOD1 silencing in ALS, we selected the widely studied SOD1 G93A (B6SJL-Tg(SOD1*G93A)1Gur/J) mouse model (*15–17, 23, 35*). This model carries the G93A mutant form of human SOD1, leading to overexpression of human mutant SOD1 protein. G93A mice have a shortened lifespan that varies depending on the background strain. Between the two background strains classically used for G93A mice, B6SJL has the shorter average lifespan of 120-140 days (*35*). We, therefore, used G93A mice with a B6SJL background. In our colony, the average lifespan is 128 ± 8 days, consistent with prior studies (*35*). Given G93A mice have a reproducibly shortened lifespan (*35*) and mouse survival is a validated endpoint in preclinical ALS studies (*35*), we selected mouse survival as our primary endpoint to evaluate di-siRNA^SOD1_123^ efficacy. 20 nmol (∼240 µg total; 120 µg in 5 µl per ventricle) of di-siRNA^SOD1_123^ or di-siRNA in the same modification pattern programmed with a non-targeting control sequence (NTC) were delivered to the CNS of pre-symptomatic 6-week-old G93A mice via intracerebroventricular (ICV) injection. Mice were then monitored for weight, clinical score, and grip strength weekly until reaching clinical endpoint (Figure 2D), defined as when mice could no longer self-right, at which time mice were euthanized (see Methods).

The median survival of naïve (untreated, n=11) and NTC-treated (n =14) mice was 128 ± 8.67 and 127 ± 8.38 days, respectively, consistent with previous studies (*15–17, 23, 35*). Treatment with di-siRNA^SOD1_123^ (n=14) nearly doubled lifespan, resulting in a median survival of 212 ± 62.9 days (Figure 2E-F, Mantel-Cox comparison of survival curves, p<0.0001). We also observed an improvement in the rate of disease progression, as measured by body weight, neurological score, and grip strength over time. The naïve and NTC-treated mice began losing weight by 15 weeks of age. By contrast, di-siRNA^SOD1_123^ treated mice stopped gaining weight at ∼20 weeks old and maintained weight until death (Figure 2G). Using an internal ALS scoring system (0 normal, 5 terminal paralysis, see Methods) to evaluate motor function, we found that naïve and NTC mice quickly progressed to near-complete hind limb paralysis by ∼20 weeks old. Initially, di-siRNA^SOD1_123^ treated mice developed motor dysfunction at the same rate as the untreated mice but began displaying a functional plateau around 15 weeks (Fig 2H) that persisted until the study endpoint. Finally, naïve and NTC-treated mice lost grip strength by week 15, whereas di-siRNA^SOD1_123^ treated mice maintained grip strength until endpoint at ∼40 weeks (Figure 2I).

SOD1 mRNA levels across the CNS were measured at the clinical endpoint (11-15 weeks post-injection for NTC mice; and 19-44 weeks post-injection for di-siRNA^SOD1_123^-treated mice). We observed significantly reduced SOD1 mRNA levels across the brain in the di-siRNA^SOD1_123^ versus NTC-treated group (50% in cortex, 55% in mid-brain, 40% in cerebellum, 40% in brainstem) (Figure 2J, two-way ANOVA with multiple comparisons, p< 0.05). There was no difference in spinal SOD1 mRNA levels between groups. These results are consistent with a higher level of di-siRNA distribution and duration of effect in mouse brain versus spinal cord following ICV injection (*25*).

### di-siRNA^SOD_123^ outperforms tofersen in extending survival in G93A mice

We next sought to benchmark the *in vivo* efficacy of di-siRNA^SOD1_123^ against the FDA-approved, SOD1-lowering ASO tofersen. To account for the different molecular sizes of di-siRNA^SOD1_123^ (∼27 kDa) and tofersen (∼7 kDa), we used a 42 nmol (∼300 µg) dose for tofersen and a 10 nmol (∼245 µg) dose for di-siRNA^SOD1_123^. To understand how SOD1 suppression impacts survival and functional outcomes relative to symptom onset, we selected two timepoints for treatment at either 6 weeks (before symptom onset) or 12 weeks (after symptom onset) of age (Figure 3A).

**Fig. 3.**
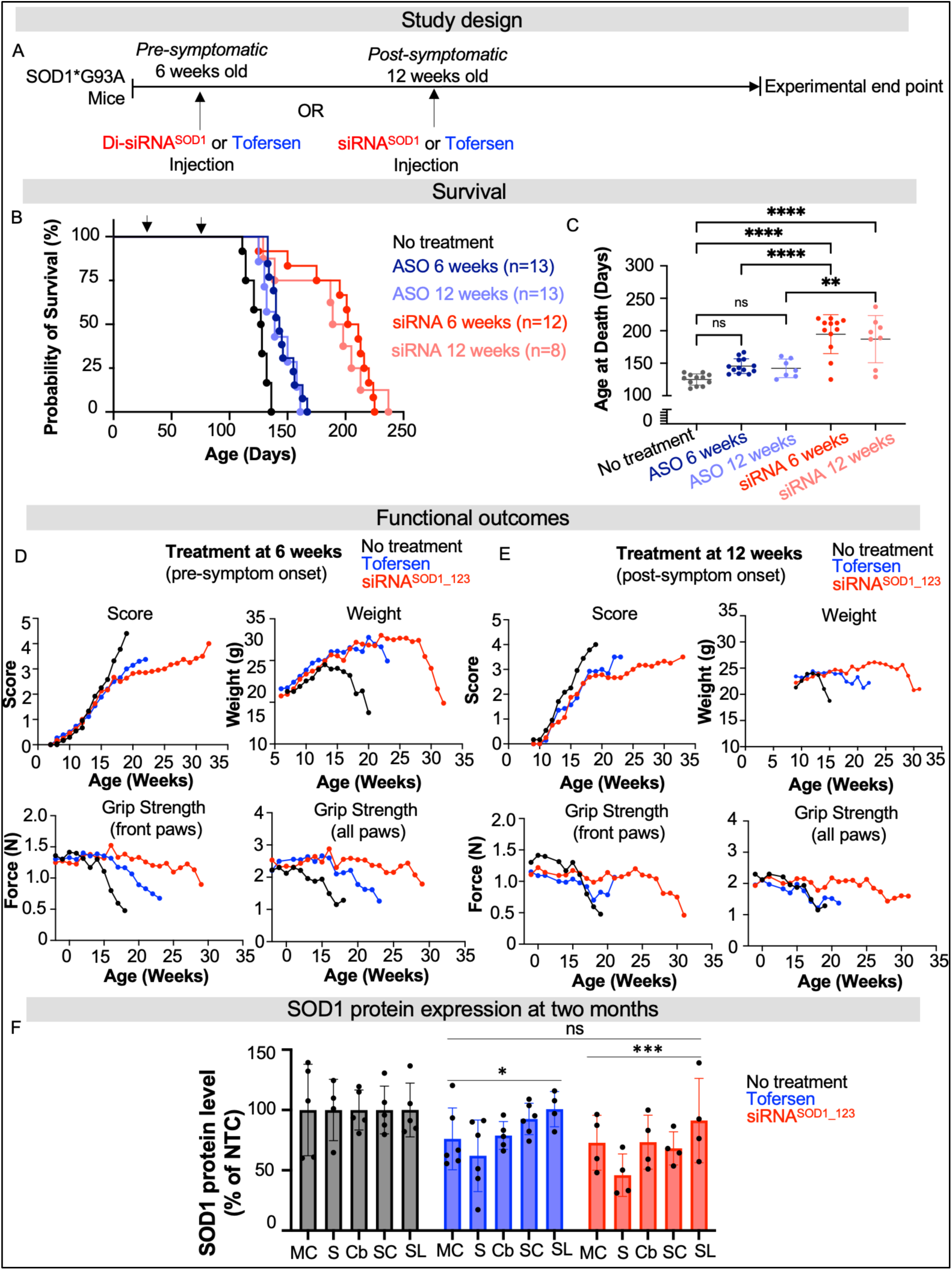
Di-siRNA^SOD_123^ showed enhanced survival compared to tofersen-treated G93A mice. (**A)** Study design. Treatment with tofersen (ASO, blue) or di-siRNA^SOD1_123^ (siRNA, red) was initiated pre- (6-weeks) or post- (12-weeks) symptom onset. (**B**) Kaplan-Meier survival curve of naïve (black), non-targeting control (NTC, gray), ASO-6 weeks (dark blue), ASO-12 weeks (light blue), siRNA-6 weeks (red) or siRNA-12 weeks (light red) G93A mice. (**C**) Plotted age at death (in days) for all groups. (**D**) Impact of NTC (black), ASO (blue) or siRNA (red) treatment at 6 weeks on ALS score, weight, and grip strength when delivered. (**E**) Impact of NTC (black), ASO (blue) or siRNA (red) treatment at 12 weeks on ALS score, weight, and grip strength. (**F**) SOD1 protein at two months post-injection across CNS regions in NTC (gray), tofersen (blue) treated, or di-siRNA^SOD1_123^ (red) treated mice. MC-medial cortex, S-striatum, T-thalamus, H-hippocampus, C-cortex, BS-brainstem, Cv-cervical spinal cord, Tc-thoracic spinal cord, L-lumbar spinal cord. * p < 0.05, ** p< 0.01, *** p < 0.001, **** p < 0.0001, Two-way ANOVA with multiple comparisons.

Treatment with tofersen at 6 weeks (n=13) or 12 weeks (n=7) led to median survivals of 143 ± 11 and 139 ± 13 days, respectively. These lifespans were not significantly different from untreated mice (median survival of 128 ± 8 days, one-way ANOVA with Tukey’s multiple comparisons test, tofersen versus untreated, 6 weeks treatment cohort p= 0.1499; 12 weeks treatment cohort p = 0.4805). By contrast, treatment with di-siRNA^SOD1_123^ at 6 (n=12) or 12 weeks (n=8) prolonged survival compared to the untreated group with median survivals of 206 ± 28 and 193 ± 34 days, respectively (Figure 3B, C) (one-way ANOVA with Tukey’s multiple comparisons test, siRNA versus untreated, 6 weeks treatment cohort p < 0.0001, 12 weeks treatment cohort p < 0.0001). Mouse survival was significantly longer with siRNA rather than tofersen treatment at 6 weeks (one-way ANOVA with Tukey’s multiple comparisons test, tofersen versus siRNA, p < 0.0001) and at 12 weeks (one-way ANOVA with Tukey’s multiple comparisons test, tofersen versus siRNA, p = 0.0023).

The impact of tofersen and di-siRNA^SOD1_123^ on disease progression in G93A mice was consistent with the observed survival data. Motor dysfunction (determined by ALS scores, see Methods) progressed at a similar rate in untreated and tofersen-treated mice (at 6 weeks), whereas di-siRNA^SOD1_123^ treatment at 6 weeks caused a plateau in disease progression (Figure 3D) similar to that observed in the prior experiment (Figure 2H). Untreated mice began losing grip strength by 15 weeks, mice treated with tofersen at 6 weeks started losing grip strength after 15 weeks, and mice treated with di-siRNA^SOD1_123^ at 6 weeks maintained grip strength for up to 30 weeks (Figure 3D). Tofersen and di-siRNA^SOD1_123^ had a similar impact on preventing G93A weight loss (Figure 3D). Treatment at 12 weeks with tofersen or di-siRNA^SOD_123^ similarly reduced progression of motor dysfunction and maintained weight compared to non-treated animals (Figure 3E). Treatment with di-siRNA^SOD_123^ maintained grip strength, unlike treatment with tofersen, compared to untreated mice (Figure 3E).

Our findings indicate that modulation of SOD1 by di-siRNA^SOD1_123^ has a more robust effect on disease progression when administered prior to the onset of symptoms and outperformed tofersen. To identify the driving factor responsible for the observed differences in outcomes between the tofersen and di-siRNA^SOD1_123^ treated groups, we first explored whether there was a difference in SOD1 reduction between groups. In a separate cohort of G93A mice treated with tofersen, di-siRNA^SOD1_123^, or NTC control at 6 weeks of age (same doses as the previous study) via ICV injection, we observed reduced SOD1 protein levels in the tofersen (10-40%) and di-siRNA^SOD1_123^ (20-60%) groups compared to NTC control at 2 months post-injection (Figure 3F, two-way ANOVA with multiple comparisons NTC vs tofersen, p < 0.05; NTC versus siRNA, p < 0.001). There was no statistically significant difference between the tofersen and siRNA^SOD1_123^ groups (Figure 3F, p > 0.05, two-way ANOVA with multiple comparisons). Two months post-injection was selected as this timepoint is when robust target silencing with di-siRNA can be observed (*25*). These results suggest that SOD1 lowering alone at an early timepoint by tofersen or siRNA may not explain survival differences between tofersen and di-siRNA^SOD1_123^ treatments over months.

### Chemical optimization of di-siRNA^SOD_123^ improves compound durability

We next investigated whether we could further improve the *in vivo* efficacy or durability of di-siRNA^SOD_123^ with additional chemical modifications. We conducted a series of *in vitro* dose-response (potency) and efficacy experiments, systematically adjusting SOD1_123 methyl content as well as guide strand 3’ end structure, to better understand which parameters impact SOD1 silencing (Figure S2). Increased methyl content in fully chemically modified siRNA has previously been shown to increase efficacy and durability (*29, 36*). In addition, a key modification of interest was the novel backbone modification, extended nucleic acid, exNA (*37*). *In vivo*, siRNA can be degraded by endo- and exonuclease, with 3’ exonuclease trimming of the guide strand being the primary degradation product detected (*38, 39*). The 3’-end exNA modification provides high resistance against 3’-exonucleases by incorporating a methylene insertion in the siRNA backbone to alter the cleavage site of these endogenous nucleases without compromising Ago2-siRNA interaction, which enables enhanced durability *in vivo* (*37*). Previously, we showed the 3’-end exNA-“UU” universal design provides an in-vivo durability benefit in the context of multiple target genes in CNS even regardless of the presence of 3’-mismatch exNA-bases on the guide strand to the target mRNAs (*37*). Thus, we assessed if the SOD1_123 sequence could also tolerate mismatches at the last two positions of the guide strand due to the UU universal sequence.

In general, we found that altering the overall structure and methyl content had a minimal impact on SOD1 silencing potency and efficacy, suggesting SOD1_123 tolerates parameter changes (Figure S2). We also demonstrate that the last two bases at the guide strand 3’ end could be modified and substituted to UU without a discernible impact on efficacy and potency (Figure S2), which is consistent with what is known about siRNAs – only positions 2-17 of the guide strand drive target complementarity; binding at the 3’ end is non-essential (*27*).

To explore chemical modifications that could optimize the *in vivo* di-siRNA performance, we then tested multiple leads *in vivo* to systemically investigate how overall 2’-O-methyl/fluoro content and 3’ end exNA modification (*37*) impacted efficacy in the CNS and survival (Figure 4A). We were first interested in exploring how enriching the siRNA 2’-O-methyl content impacted in vivo silencing efficacy and survival (Figure 4B,C). *In vitro*, we found that 2’-O-methyl-rich SOD1_123 had a significantly lower IC50 value than a 2’-O-methyl-balanced SOD1_123 (passive uptake, 8.17 nM versus 46.3 nM, respectively) (Figure 4D). We injected 20 nmol of the original 2’-O-methyl-balanced di-siRNA^SOD1_123^ (Figure 4B), 2’-O-methyl-rich di-siRNA^SOD1_123^ (Figure 4C), or NTC into G93A mice to compare *in vivo* efficacy. At 2 months post-injection, both balanced and methyl-rich di-siRNA^SOD1_123^ significantly lowered SOD1 protein compared to NTC (balanced 10-50%, methyl-rich 20-60%, two-way ANOVA with multiple comparisons, NTC versus balanced, p< 0.001; NTC versus methyl-rich, p < 0.0001, balanced versus methyl-rich, p < 0.05), with the methyl-rich pattern improving knockdown efficacy over the methyl-balanced pattern (Figure 4E, M, medial cortex; S striatum, B, brainstem, C, cervical spine, L, lumbar spine). To determine if the improved silencing efficacy of 2’-O-methyl-rich di-siRNA^SOD1_123^ translated to improved survival in mice, we delivered 2’-O-methyl-rich di-siRNA^SOD1_123^, 2’-O-methyl-balanced di-siRNA^SOD1_123^, or NTC to 6-week-old G93A mice via ICV injection and monitored survival. Although both treatments prolonged lifespan compared to control mice, methyl-balanced di-siRNA^SOD1_123^ outperformed methyl-rich di-siRNA^SOD1_123^ by ∼20 days (Figure 4F). These results suggest that SOD1 protein levels *in vivo* at an early time point post-injection (i.e., 2 months) is not predictive of long-term survival in G93A mice. For subsequent experiments, we relied on survival results in mice to determine *in vivo* di-siRNA efficacy.

**Fig 4.**
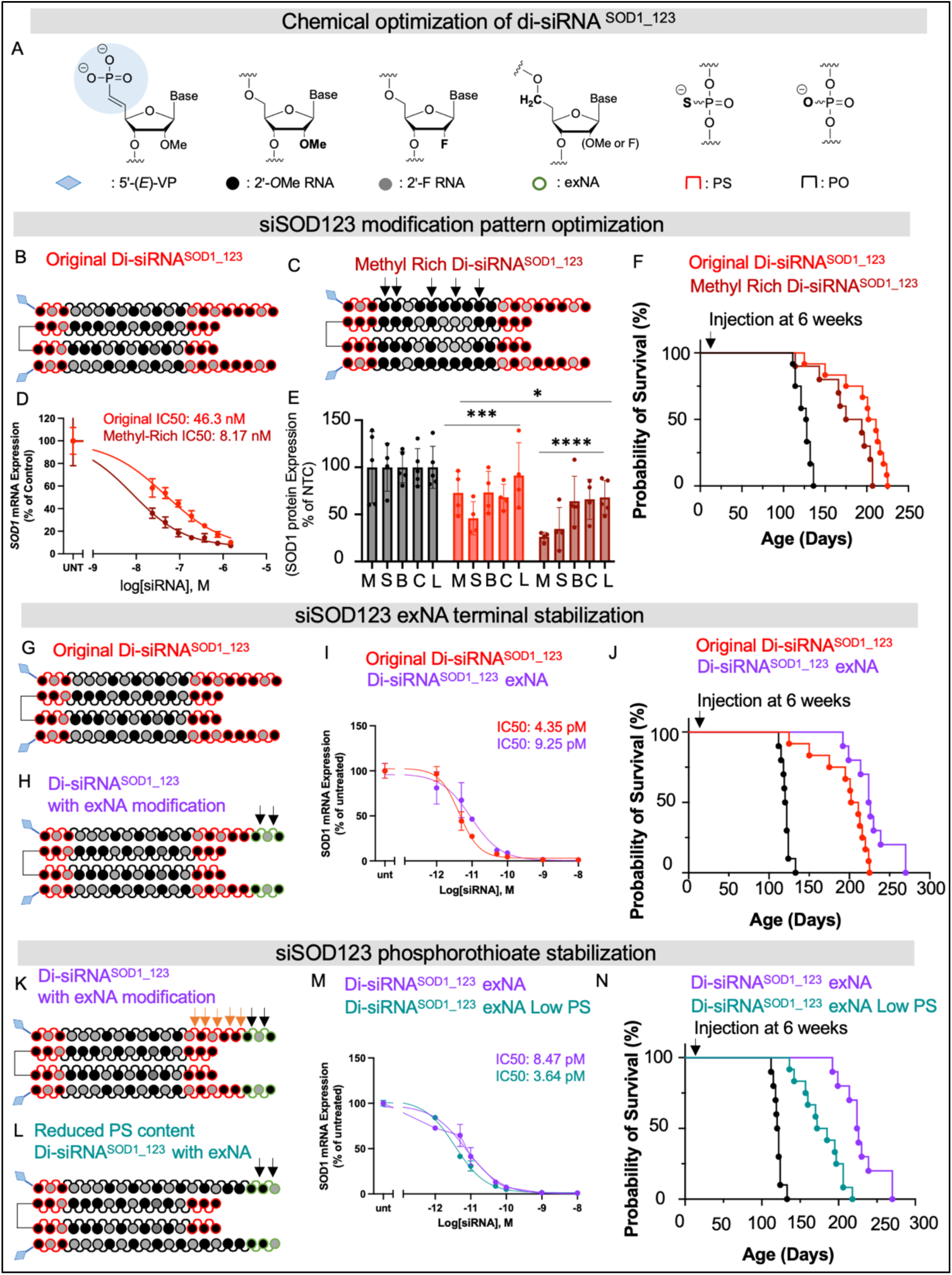
Chemical optimization of di-siRNA^SOD_123^ enhances the impact on survival. (**A**) chemical modifications used to optimize di-siRNA^SOD_123^. (**B**) chemical modifications and structure of original di-siRNA^SOD_123^ (**C)** chemical modifications and structure of methyl-rich di-siRNA^SOD_123^ (**D**) *In vitro* dose response of original (red) or methyl-rich (maroon) di-siRNA^SOD_123^ (**E**) *in vivo* protein silencing of original or methyl-rich siRNA at two months post-injection. (**F)** Kaplan-Meier survival curve comparing G93A mice treated with the original or methyl-rich di-siRNA^SOD_123^ or NTC control. (**G**) Chemical modifications and structure of original di-siRNA^SOD_123^. (**H**) Chemical modifications and structure of di-siRNA^SOD_123^ exNA (exNA noted with black arrows). (**I**) 7-point dose-response curves of original (red) or exNA (purple) di-siRNA^SOD_123^ (**J**) Kaplan-Meier survival curve comparing G93A mice treated with the original or exNA di-siRNA^SOD_123^ or NTC control. (**K**) Chemical modifications and structure of exNA di-siRNA^SOD_123^ (PS noted with orange arrows, exNA with black arrows) (**L**) Chemical modifications and structure of exNA di-siRNA^SOD_123^ with reduced phosphorothioate content (exNA noted with black arrows). (**M**) 7-point dose-response curves of exNA di-siRNA^SOD_123^ high PS (purple) or exNA di-siRNA^SOD_123^ low PS (teal). (**N)** Kaplan-Meier survival curve comparing G93A mice treated with exNA di-siRNA^SOD_123^ high PS (purple), exNA di-siRNA^SOD_123^ low PS, or standard NTC control. Survival studies used single injection at 6 weeks.

We synthesized the original 2’-O-methyl-balanced (Figure 4G) or a 2’-O-methyl-balanced di-siRNA^SOD1_123^ compound with 3’ end exNA modification (Figure 4H). *In vitro*, di-siRNA^SOD1_123^ with 3’ end exNA exhibited similar potency (lipid-mediated uptake, IC50 9.25 pM) to the original 2’-O-methyl-balanced di-siRNA^SOD1_123^ (lipid-mediated uptake, IC50 4.35 pM) (Figure 4I). However, 20 nmol di-siRNA^SOD1_123^ with 3’ end exNA delivered to 6-week-old G93A mice via ICV injection extended survival past 250 days, outperforming the original di-siRNA^SOD1_123^ (Figure 4J). The same cohort of untreated and NTC-treated mice (Figure 2E) were used as the control arm for all siRNA optimization studies to minimize the number of animals used.

Finally, with the improved survival observed following di-siRNA^SOD1_123^ exNA treatment, we sought to understand if we could lower the phosphorothioate (PS) content while maintaining efficacy and durability. PS is a gold standard for stabilizing therapeutic RNA modalities including siRNA that supports long plasma circulation and durability in vivo (*32, 40, 41*). However, PS, especially in an ASO context, can be associated with toxicity *in vivo* (*40*). Indeed, all clinically advanced ASO compounds have a mixed backbone modification pattern where the number of PS modifications is reduced (*28, 42*). With the presence of exNAs in the siRNA, we hypothesized that it may be possible to remove some of the PS to improve safety while maintaining siRNA efficacy and durability. We synthesized a version of di-siRNA^SOD1_123^ exNA with reduced PS content near the 3’ end of the guide strand (Figure K,L). *In vitro,* di-siRNA^SOD1_123^ exNA with low PS content exhibited similar potency (IC50 3.64 pM) to di-siRNA^SOD1_123^ exNA (IC50 8.47 pM) (Figure 4M); however, G93A mice survival was significantly lower following treatment with the low PS compound at 6 weeks (Figure 4N). Taken together, these studies identify 2’-O-methyl-balanced di-siRNA^SOD1_123^ exNA with 7 PS modifications as the top compound for further biological and neuropathological studies.

### Repeat injections of optimized di-siRNA^SOD_123^ further extend G93A mouse survival

Repeat injections are a common practice in medicine to sustain target suppression. To determine whether multiple injections of 2’-O-methyl-balanced di-siRNA^SOD1_123^ exNA could further extend survival, we delivered our lead compound at 20 nmol/10 µL (240 µg) to G93A mice via ICV injection at 6 weeks old (single injection) or at 6, 14, and 22 weeks old (repeat injection, Figure 5A) compared to a no-treatment G93A control arm. Multiple injections of 2’-O-methyl-balanced di-siRNA^SOD1_123^ exNA extended mouse survival past 340 days, with median survival being 281 ± 41 days (Figure 5B). This lifespan is significantly longer than the single dose of 2’-O-methyl-balanced di-siRNA^SOD1_123^ with or without exNA. In both the single-dose cohort and multiple-dose cohort, di-siRNA^SOD1_123^ exNA-treated mice maintained weight (Figure 5C) and exhibited slower progression of motor dysfunction compared to untreated mice (Figure 5D). To ensure that the observed impact on these outcomes was due to SOD1 lowering, we delivered 2’-O-methyl-balanced di-siRNA^SOD1_123^ exNA at 20 nmol/10µL (240 µg) into a separate cohort of G93A mice via ICV injection and measured SOD1 mRNA in cortex and cervical spinal cord at two months post-injection. As controls, we measured SOD1 mRNA levels in untreated G93A mice, naive wild-type (WT) mice, and mice expressing WT human SOD1. Two months post-injection, the 2’-O-methyl-balanced di-siRNA^SOD1_123^ exNA demonstrated 60-75% SOD1 mRNA reduction in both brain and spinal cord compared to untreated G93A mice (Figure 5E,F, one-way ANOVA with Tukey’s multiple comparison, G93A untreated versus siRNA treated, cortex p < 0.001, cervical spinal cord p < 0.0001).

**Fig. 5.**
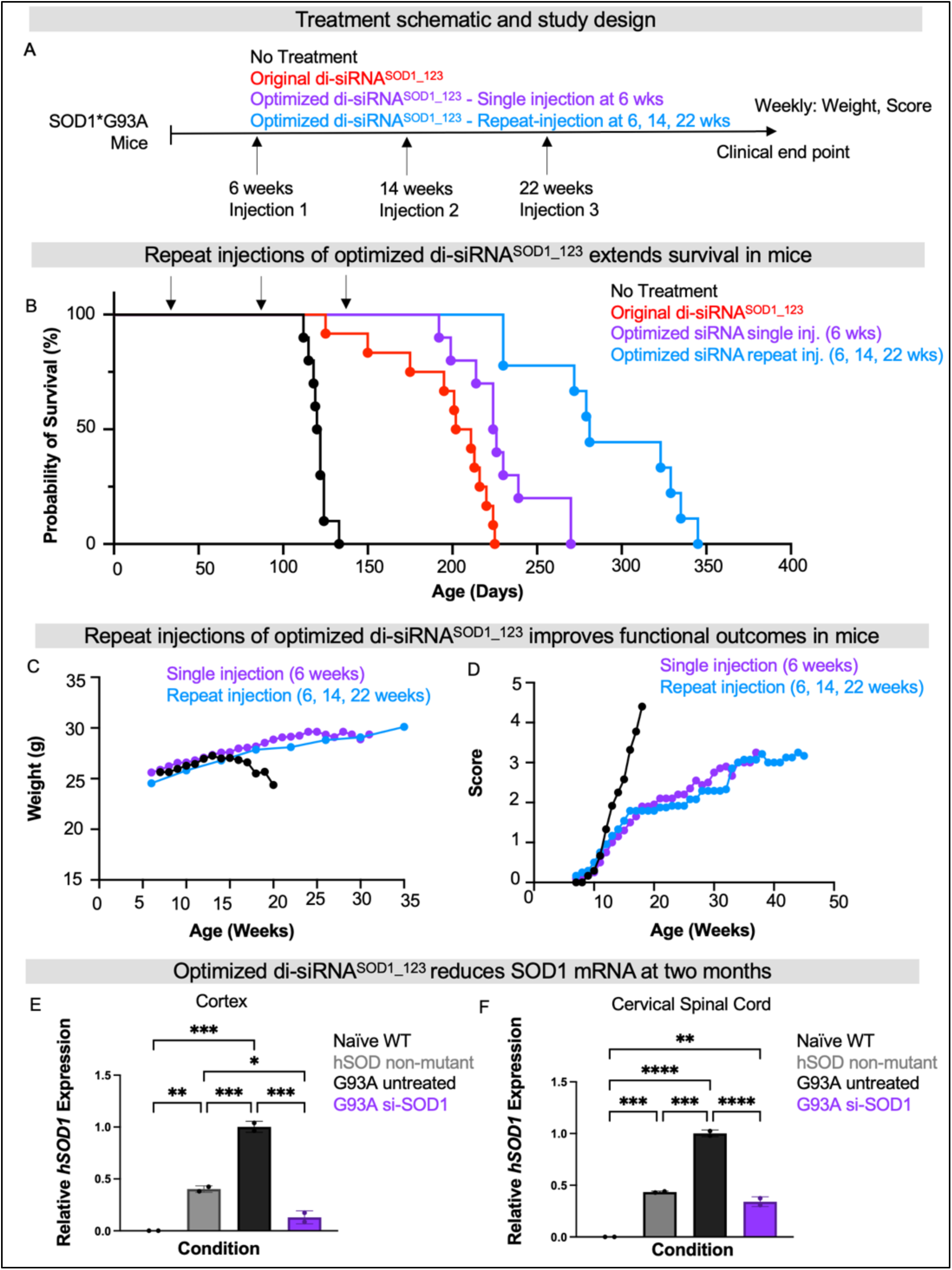
Repeat injections of di-siRNA^SOD_123^ further enhance survival. (**A)** Study design: no treatment (black), original di-siRNA^SOD_123^ (red), single injection of exNA di-siRNA^SOD_123^ (siRNA, purple) or repeat injections (6, 14, 22 weeks) of exNA di-siRNA^SOD_123^ (light blue) arms. Treatment was initiated at 6-weeks of age (pre-symptom onset). (**B**) Kaplan-Meier survival curve comparing G93A mice with no treatment, single injection of the original di-siRNA^SOD_123^ treatment or exNA di-siRNA^SOD_123^ treatment, or repeat injection of exNA di-siRNA^SOD_123^. (**C**) Weight and (**D**) clinical scores for each study arm tracked until clinical endpoint. (**E**) SOD1 cortical mRNA levels (*hSOD1*, human SOD1) in naïve WT, humanized non-mutant SOD1 mice, G93A untreated mice, or the exNA di-siRNA^SOD_123^ (single injection) arm at clinical endpoint. (**F**) SOD1 cervical mRNA levels (*hSOD1*, human SOD1) in naïve WT, humanized non-mutant SOD1 mice, G93A untreated mice, or the exNA di-siRNA^SOD_123^ (single injection) arm at clinical endpoint. Tissue collection was performed and evaluated for SOD1 mRNA levels at the following time points: 174, 145, 70-100, or 230 days, respectively.

### Repeat injections of optimized di-siRNA^SOD_123^ maintain synaptic structure and prevent neuron loss and in the motor cortex and spinal cord

Beyond survival, it is crucial to understand to what extent SOD1 lowering by our lead compound protects against cortical and peripheral neuron loss and dysfunction – hallmarks of ALS. We conducted a panel of neuropathological assessments on and the di-siRNA^SOD1_123^ exNA-treated G93A mice from the prior study with naive WT mice, transgenic mice with the non-mutant human SOD1, untreated mutant G93A human SOD1 as controls (Figure 6A). To assess neuron count and health, the cortex (Figure 6B) and spinal cord (Figure 6C) of mice were evaluated with Nissl stain. Untreated G93A mice used for this analysis were 70-100 days old. At this timepoint, we note that neuron loss is not expected in the cortex or spinal cord, but additional synaptic or apoptotic pathways may be involved at this age. We controlled for this by including transgenic non-mutant and wild-type mice as additional groups. We observed a 20-fold reduction in pyknotic neurons in di-siRNA^SOD1_123^ exNA-treated G93A mice (Figure 6D), approaching WT levels. We found that treated G93A mice showed no difference in total cortical neuron count (Figure 6E) or lumbar spinal cortex count (Figure 6F) compared to untreated G93A mice. Using the same control groups (Figure 7A), we next evaluated synaptic integrity in the motor cortex (Figure 7B) and spinal cord (Figure 7C). The most profound histological changes we saw were a significant increase in PSD-95 positive synapses in the motor cortex (Layer V) (Figure 7D) and anterior horn of the lumbar spinal cord (Figure 7E) between treated and untreated G93A mice. Moreover, we found that di-siRNA^SOD1_123^ exNA treatment almost fully reversed a significant decrease in the synapse marker, synaptophysin, in the motor cortex (Figure 7F) and lumbar spinal cord (Figure 7G) of G93A mice.

**Figure 6.**
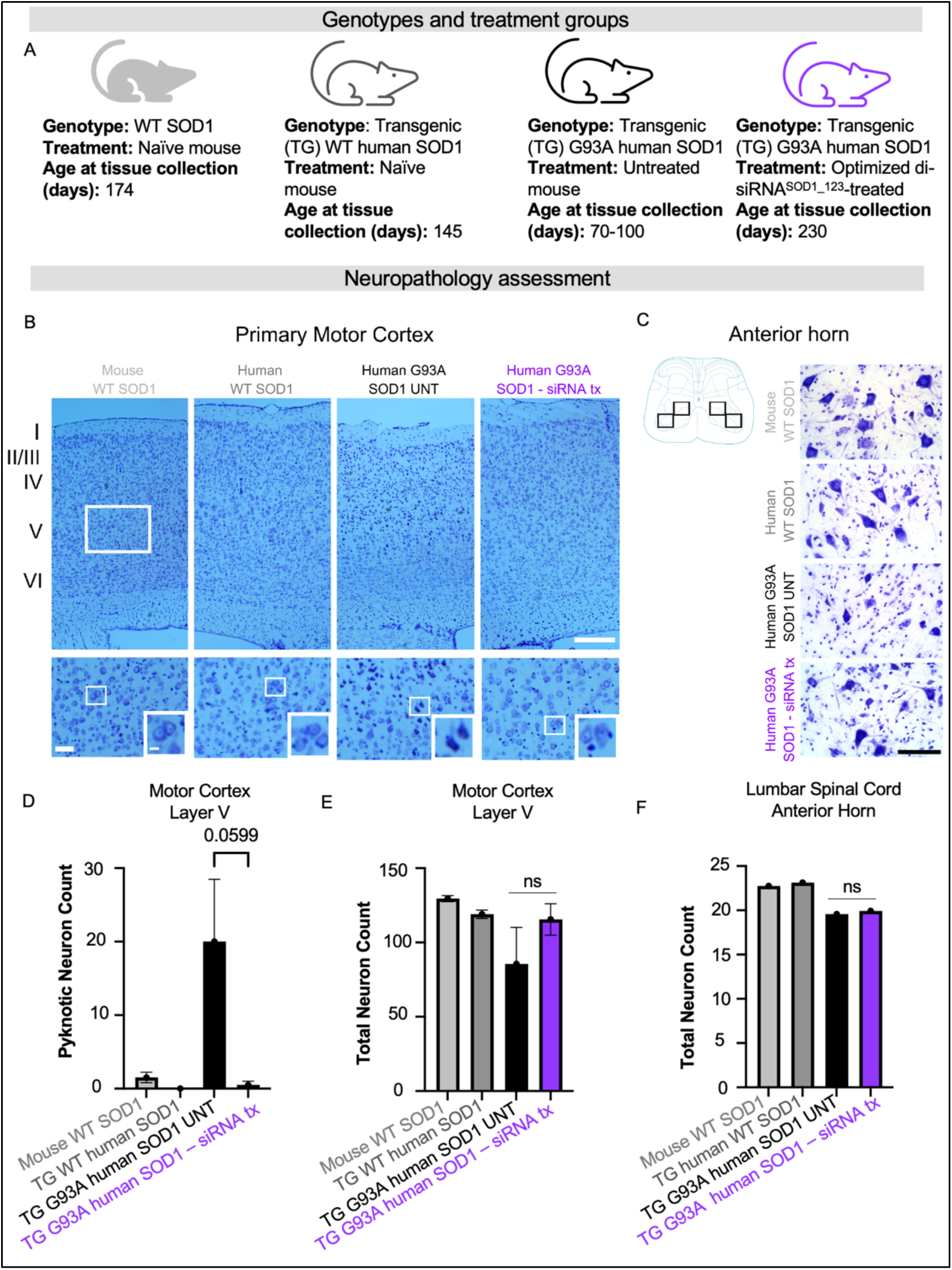
SOD1 knockdown with di-siRNA decreases number of apoptotic dark neurons in layer V of primary motor cortex in G93A mice. **(A)** Genotypes and treatment groups used to assess neuropathology (**B**) Representative images of primary motor cortex, rectangles indicate the approximate region of interest (ROI) in layer V used for quantitative analyses. Scale bar = 250 µm (low power magnification), Scale bar=50 µm (high power magnification). **(C)** Cartoon and representative images showing ROIs used for quantification of pyramidal neurons. Scale bar=100 µm. **(D)** total number of Nissl-stained cortical neurons showed no statistical difference between groups. **(E)** dark neurons were significantly higher in number in the hSOD1^G93A^ mice without treatment compared with other groups. **(F)** Number of pyramidal neurons counted within four ROIs in the anterior horn (cartoon) (average from two sections per animal). Data in the bar graphs are shown as mean ±SEM n=2 per group.

**Figure 7:**
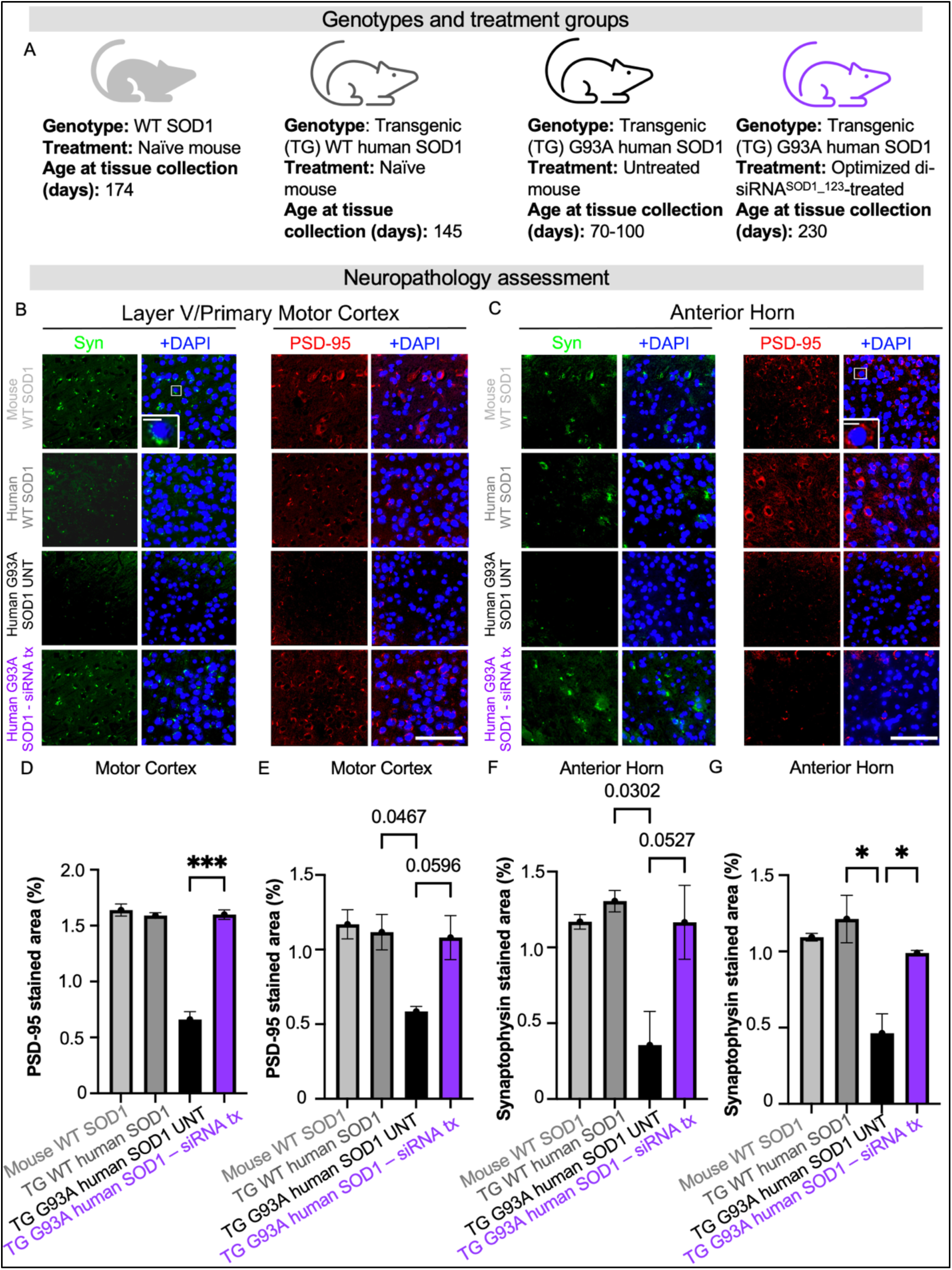
SOD1 silencing with di-siRNA rescued loss of synaptic markers in G93A mice. **(A)** Genotypes and treatment groups used to assess neuropathology. **(B)** Representative images and insets showing presynaptic marker synaptophysin and postsynaptic marker PSD-95 staining signal from layer V of primary motor cortex. **(C)** Representative images and insets showing synaptophysin and PSD-95 staining signal from anterior horn of the lumbar spinal cord**. (D)** Quantification of PSD-95 in layer V of primary motor cortex. **(E)** Quantification of PSD-95 in anterior horn. **(F)** Quantification of synaptophysin in layer V of primary motor cortex (**G**) Quantification of synaptophysin in the anterior horn. Scale bar=100 µm, Scale bar=10 µm (inset). Data in the bar graphs are shown as mean ±SEM n=2 per group.

To determine if SOD1 reduction by di-siRNA^SOD1_123^ exNA in G93A mice impacted neurogliosis, we evaluated astrocyte and microglial morphology in the cortex and spinal cord. In the cortex, there was no evidence of either astrogliosis (Figure S3A,C) or microgliosis (Figure S4A,C) between wild-type, transgenic non-mutant or transgenic mutant untreated mice, while both phenomena were enhanced in di-siRNA^SOD1_123^ exNA treated G93A mice. By contrast, in the lumbar spinal cord, untreated G93A mice exhibit both astrogliosis (Figure S3B,D) and microgliosis (Figure S4B,D) compared to wild-type or non-mutant transgenic mice that was unaffected by the di-siRNA^SOD1_123^ exNA treatment.

Taken together, these immunohistochemistry findings suggest that di-siRNA^SOD1_123^ exNA treatment maintains synapse structure and protects against the synapse loss observed in untreated ALS mice. Concurrently, di-siRNA^SOD1_123^ exNA provokes astroglial and microglial activation in the motor cortex but not spinal cord.

### Repeat injections of di-siRNA^SOD_123^ exNA preserve neuromuscular junction innervation

In patients with ALS and in the G93A mouse model, death of the innervating motor neurons triggers denervation of the NMJ, leading to weakness and paralysis. We, therefore, sought to understand how di-siRNA^SOD1_123^ exNA impacted the structural integrity of the NMJ in G93A mice. Using a single mouse from the naïve WT, non-mutant humanized SOD1, untreated G93A, and di-siRNA^SOD1_123^ exNA-treated G93A groups from the prior experiment, we assessed NMJ integrity in the gastrocnemius muscle, with post-synaptic NMJ stained with alpha bungarotoxin and the pre-synaptic NMJ stained with synaptophysin (Figure 8A). As expected, the NMJs were intact (measured by the percent of pre- and post-synapse co-localization in the naïve WT mouse) in the naïve and WT transgenic mouse. Substantial denervation was observed in the untreated G93A mouse. Strikingly, the structural integrity of the NMJ was fully rescued to WT levels in the di-siRNA^SOD1_123^ exNA-treated G93A mouse (Figure 8B). These results suggest that di-siRNA^SOD1_123^ exNA treatment protects the NMJ in G93A mice.

**Fig. 8.**
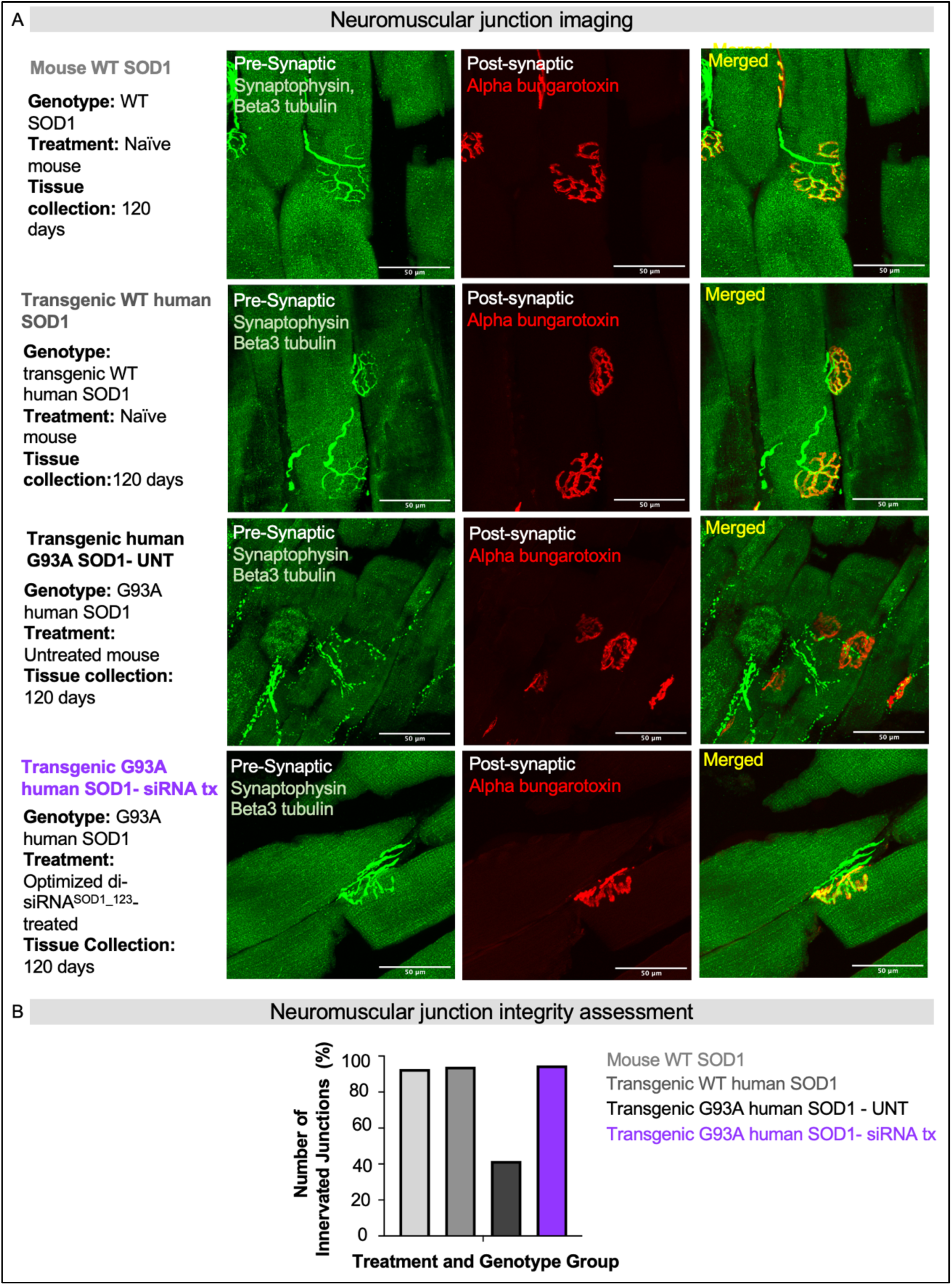
exNA di-siRNA^SOD_123^ prevents neuromuscular junction destruction. (**A**) Representative images from gastrocnemius slices co-stained with synaptophysin and beta-tubulin (green, pre-synapse) or alpha-bungarotoxin (red, post-synapse) to mark the neuromuscular junction (NMJ) (Merged, yellow). Scale bar = 50 µm. (**B**) Quantified NMJ innervation (percentage, %) for each treatment and genotype group. naïve WT, n=1; humanized non-mutant SOD1 mice, n=1; G93A untreated mice, n=1; the SOD1 siRNA-treated G93A, n=1.

## DISCUSSION

SOD1 ALS is a uniformly lethal disorder with limited treatment options. Currently, the only FDA-approved therapy specifically for SOD1 ALS is tofersen, a GapmeR ASO that lowers SOD1 levels (*21*). However, tofersen shows modest clinical efficacy. Other modalities that may provide alternative PK/PD properties could ultimately further improve outcomes for patients. Here, we designed, screened, validated, and optimized a chemically stabilized siRNA that successfully reduces SOD1 in the CNS of G93A ALS mice after a single injection, and significantly outperformed tofersen in enhancing the survival of G93A mice. Repeat injections of this chemically-optimized siRNA enhanced lifespan even further, prevented neuron loss in the motor cortex and spinal cord, and maintained synaptic structural integrity in the motor cortex, lumbar spinal cord, and at the neuromuscular junction. This work identifies a pre-clinical lead for ALS patients with familial SOD1 ALS.

### Comparing tofersen and di-siRNA

Tofersen is the key benchmark for new approaches to SOD1 suppression (*18, 21, 33*). In this study, tofersen and di-siRNA^SOD1_123^ showed similar effects on SOD1 protein reduction at two months post-treatment but profoundly different effects on long-term survival, with di-siRNA^SOD1_123^ outperforming tofersen.

The observed differences in survival between di-siRNA and tofersen treatment may be due to differences in mechanisms of action and durability/stability of each oligonucleotide. ASOs first engage with an mRNA target, after which the complex is recognized by RNAse-H, resulting in mRNA cleavage. The ASO can potentially recycle to different mRNA targets but only a limited number of times (*43*). For siRNAs, the first step is loading into the RNA-induced silencing complex (RISC), which then recognizes and cleaves the target mRNA (*44*). Upon cleavage, RISC is recycled efficiently with hundreds of loaded complexes per cell, thereby maintaining complete target gene modulation for a longer time. Thus, although both modalities show robust efficacy in the CNS, the duration of effect of optimized siRNAs tends to surpass that of ASOs (*43*).

For both ASOs and siRNAs, endosomal entrapment serves as an intracellular drug depot enabling slow release into the cytoplasm and protracted efficacy over several months. Chemical stabilization of oligonucleotides is crucial to prevent degradation in endosomal/lysosomal environments. Nusinersen, a steric-blocking ASO (*45, 46*), achieves high stability through methoxyethyl modification of all sugars (*47*). However, GapmeR ASOs like tofersen, requiring an unmodified DNA gap for RNaseH recognition, face stability limitations inside endosomes. The increased chemical stabilization of siRNAs, compared to tofersen, likely contributes to their longer-lasting effects.

Here we show that a SOD1-targeting siRNA improved disease phenotypes in ALS mice compared to tofersen or less optimized siRNA. While higher dosing of tofersen might replicate a similar effect, the current dosing regimen is constrained by safety concerns, with 100 mg required every two weeks for three doses followed by 100 mg re-injections every 28 days, considered the maximally tolerated safe dose (*21*). Tofersen’s therapeutic index is relatively narrow, with CNS toxicity observed in other ASO trials (e.g., Phase 3 tominersen trial for Huntington’s disease (*48*). Potent and durable compounds like siRNAs offer the potential for reduced dosing frequency and more significant disease impact, potentially preventing or halting disease progression.

Here we utilized ICV injection, commonly used in rodent models, including ASOs (*49*). In patients, intrathecal injection is a more likely route of administration. In large animals, both intrathecal and ICV administrations delivered siRNA to the spinal cord and cortex, key areas in ALS (*50*), suggesting di-siRNA^SOD1_123^ may also work as an intrathecal injection for straightforward clinical translation.

### Alternative approaches to SOD1 suppression

SOD1 lowering can be pan-allele (non-mutant and mutant) or mutant allele-specific. Tofersen and SOD1_123 characterized here are both pan-allele silencing. In preclinical models and one human study, non-selective (e.g., ASOs) SOD1 lowering has shown to be a successful strategy to improve ALS-related outcomes (*17, 51, 52*). *In vivo*, an siRNA with ASO-conjugate intrathecally delivered extended G93A SOD1 ALS mouse survival until ∼240 days highlighting the benefit of siRNA to target SOD1 in the setting of ALS (*23*). G93A survival until 240 days (at the dose used here) is comparable to the extension in life observed with di-siRNA^SOD1_123^. Once the di-siRNA^SOD1_123^ was modified with exNA, we were able to extend the durability of the SOD1 lowering treatment with G93A survival persisting until ∼260 days at the same dose without need for reinjection.

Collectively, this work suggests that using oligonucleotides for non-specific allele suppression holds promise for clinical benefits. However, establishing the ideal SOD1 reduction level for both wild-type and mutant alleles, along with ensuring safety, requires clinical evaluation. Beyond siRNA and ASOs, other modalities including small molecules, splice modulators, CRISPR gene editing, and adeno-associated virus (AAV) based therapies, have been considered for non-allele and allele selective suppression of SOD1 in ALS (*53*). In preclinical models, small molecules have been used to lower SOD1 levels or inhibit SOD1 aggregation *in vitro* (*54, 55*). SOD1 lowering with short-hairpin RNA (shRNA) delivered by AAV or a small molecule protected motor neurons as well as improved neurodegenerative phenotypes (*16, 56*). In patients, SOD1 suppression with approaches other than tofersen has been evaluated with varying success.

The small molecule pyrimethamine was studied in an open-label, dose-finding study of 32 patients and showed significant lowering of CSF SOD1 levels. The drug was tolerated at doses <100 mg and but in a pilot study, no major impact on disease progression, measured by quality of life and functional outcomes, was observed (*57*). In a pilot trial with two ALS patients, AAV-delivered synthetic miRNA temporarily stabilized or improved ALS scores and functions, reducing SOD1 levels in the spinal cord (*51*). However, one patient experienced transient liver dysfunction and sensory radiculitis due to immunoreactivity to viral capsids. Immunosuppressive drugs prevented these adverse events in the second patient, indicating safe SOD1 reduction using alternative methods.

### Di-siRNA SOD1- lowering on G93A neuroinflammation

Understanding the impact of SOD1 reduction by siRNA on CNS inflammation is crucial. Our study revealed that while SOD1 lowering with di-siRNA influenced neuroinflammation in G93A mice, it had differing effects in the cortex and spinal cord. In the cortex, di-siRNA triggered astrogliosis and microgliosis, possibly due to proximity to the intracerebroventricular injection site. However, in the spinal cord, di-siRNA attenuated disease-induced microgliosis, suggesting a region-specific response. This disparity could be due to variations in astrocyte and microglia composition or distribution between the cortex and spinal cord. Monitoring the impact of di-siRNA^SOD1_123^ on cortical and spinal neuroinflammation in patients will be crucial if it progresses as a clinical candidate, considering regional differences in the CNS.

### Modeling ALS in G93A mice

Considering the timing of SOD1 lowering intervention in ALS, it’s vital to recognize early and late disease signs. Our study in G93A mice showed that siRNA treatment rescued synapse loss in the cortex and spinal cord, occurring before neuron loss. This suggests that synapse integrity decline may precede neuron loss, indicating a potential earlier clinical endpoint for treatment efficacy assessment. Importantly, our findings suggest that both pre-symptomatic and symptomatic siRNA treatment improved survival and clinical scores, indicating the potential benefit of intervention even after symptom onset.

### Clinical translation of future SOD1 targeting compounds

This study revealed that *in vitro* silencing efficacy does not reliably predict in vivo efficacy for di-siRNA. Additionally, the *in vivo* efficacy of either tofersen or di-siRNA at 2 months did not correlate with survival outcomes. The similarity in mRNA or protein level silencing did not explain the significant survival differences observed. *In vitro* efficacy relies on factors like siRNA internalization and target binding over short periods (72-hours), while *in vivo* efficacy involves more complex processes over months, such as CSF circulation and neuronal uptake. Thus, factors like size, stability, and durability become critical for *in vivo* evaluation. Clinical outcomes like survival should guide preclinical compound assessment, although biochemical readouts like CSF Nfl or SOD1 levels are important. Care should be taken in predicting clinical or behavioral outcomes solely based on biochemical readouts, although such readouts are important.

Because tofersen is already FDA-approved, any clinical trial design for other compounds will require a demonstration of superiority versus tofersen. Considering the high dose of tofersen currently used and relatively limited therapeutic index, a combinatorial treatment might be hard to execute safely. Thus, innovative and dynamic clinical trial design would need to be considered to advance novel therapeutic options for SOD1 ALS.

### Nucleic acid therapeutics beyond SOD1 ALS

Advancing ALS treatment involves addressing various ALS-causing mutations. Di-siRNA and similar configurations, including C16 (*24*), offer versatility, as their structure allows for easy reprogramming to target different SOD1 mutations or other common ALS genes like C9orf72 (*58–60*), FUS (*61*) and TARDBP (*62*). These chemically stabilized siRNAs are also being explored for other CNS diseases like Huntington’s and Alzheimer’s, by groups including Arrowhead (NCT05949294) and Alnylam (NCT05231785) (*63*). Improvements in delivery, durability, and distribution are enhancing the efficacy of CNS-targeted siRNAs, with several clinical programs underway.

## Conclusions

In conclusion, the development of nucleic acid therapeutics, particularly in the CNS, offers new hope for treating neurodegenerative disorders. Our study highlights the significant lifespan extension and improved neurodegenerative outcomes in ALS through SOD1 reduction with chemically stabilized siRNA. This represents a promising, durable, and more effective therapy for SOD1 ALS patients.

## MATERIALS AND METHODS

### Oligonucleotide synthesis

Oligonucleotides were synthesized by solid-phase synthesis using the phosphoramidite method on a MerMade12 (Biosearch Technologies, Novato, CA) or on a Dr Oligo 48 (Biolytic, Fremont, CA). Phosphoramidites used include 2’-Fluoro and 2’-O-Methyl modified phosphoramidites with standard protecting groups, 5’-(E)-Vinyl tetraphosphonate (pivaloyloxymethyl) 2’-O-methyl-uridine 3’-CE phosphoramidite, and exNA phosphoramidites (2’-OMe ex-Ur, 2’-Fluoro ex-Ur) purchase from ChemGenes, Wilmington, MA, and Hongene Biotech, Union City, CA. Phosphoramidites were prepared at 0.1 M in anhydrous acetonitrile (ACN) or 15% dimethylformamide in ACN. 5-(Benzylthio)-1H-tetrazole (BTT) was used as the activation reagent at 0.25 M. Coupling time was 4 min with 10 eq. Detritylations were performed using 3% trichloroacetic acid in dichloromethane. Capping reagents used were CAP A (20% N-methylimidazole in ACN) and CAP B (20% acetic anhydride and 30% 2,6-lutidine in ACN).

Phosphite oxidation to convert to phosphate or phosphorothioate was performed with 0.05 M iodine in pyridine-H2O (9:1, v/v) or 0.1 M solution of 3-[(dimethylaminomethylene)amino]-3H-1,2,4-dithiazole-5-thione (DDTT) in pyridine for 3 min, respectively. All reagents were purchased from Chemgenes. Cholesterol conjugated oligonucleotides were synthesized on a 500Å LCAA-CPG solid support, Tetraethyleneglycol cholesterol moiety was the functionalized group joined by a succinate linker (Chemgenes). Divalent oligonucleotides (DIO) were synthesized on a 500Å long-chain alkyl amine (LCAA) controlled pore glass (CPG) solid support functionalized via succinyl linker with a glycerol-tetraethyleneglycol linker (Hongene Biotech). Unconjugated oligonucleotides were synthesized on a 500Å LCAA-CPG solid support functionalized with a Unylinker terminus (Chemgenes).

### Deprotection and Purification of Oligonucleotides for Sequence Screening

Oligonucleotides with or without cholesterol conjugate used for in vitro experiments were cleaved and deprotected on-column with Ammonia gas (Airgas Specialty Gases) and a modified on-column ethanol precipitation protocol was used for purification.

### Deprotection and Purification of Oligonucleotides for in vivo Experiments

5’-(E)-Vinyl-phosphonate containing oligonucleotides were cleaved and deprotected with 3% diethylamine in ammonium hydroxide, for 20 h at 35℃ with slight agitation. Unconjugated and DIO oligonucleotides were deprotected with 1:1 ammonium hydroxide and aqueous monomethylamine, for 2 h at 25℃ with slight agitation. The controlled pore glass was subsequently filtered and rinsed with 30 mL of 5% ACN in water and dried overnight by centrifugal vacuum concentration. Purifications were performed on an Agilent 1290 Infinity II HPLC system using Source 15Q anion exchange resin (Cytiva, Marlborough, MA). The loading solution was 20 mM sodium acetate in 10% ACN in water, and elution solution was the loading solution with 1M sodium bromide, using a linear gradient from 30 to 70% in 40 min at 50°C. Pure fractions were combined and desalted by size exclusion with Sephadex G-25 (Cytiva).

Purity and identity of fractions and pure oligonucleotides were confirmed by IP-RP LC/MS on an Agilent 6530 Accurate-mass Q-TOF.

### *In vitro* and *in vivo* preparation of oligonucleotides

siRNA single strands were mixed at a 1:1 ratio in water. To anneal the duplex, the single strands were heated to 95C for 15 minutes and allowed to cool to room temperature for at least one hour. The duplexes were run on a 20% TBE gel compared to single arm oligonucleotide controls to assure duplexing quality. In vitro siRNA (cholesterol conjugated) were prepared in water. Duplexes for in vivo injection (divalent siRNA) were lyophilized using a speed vacuum and resuspended in PBS to a final concentration of 20 nmol of active siRNA guide strand per 10 µL PBS per mouse (240 µg total).

### Cell Culture (HeLa, LLC-MK2, and SH-SY5Y cells)

HeLa (ATCC, #CCL-2) and LLC-MK2 (ATCC, #CCL-7) cells were maintained in Dulbecco′s Modified Eagle′s Medium (DMEM) (Cellgro, #10-013CV). All media was supplemented with 9% fetal bovine serum (FBS) (Gibco, #26140), and all cells were grown at 37°C and 5% CO2. Cells were split every 3 to 7 days and discarded after 15 passages.

### Passive uptake of oligonucleotides

HeLa cells (ATCC, #CCL-2) were plated in DMEM (Cellgro, #10-013CV) containing 6% FBS (Gibco, #26140) at 10,000 cells per well in 96-well tissue culture plates. Cholesterol-conjugated siRNA was diluted in OptiMEM (Gibco, #31985-088) and added to cells, resulting in 3% FBS final. Cells were incubated for 72 h at 37°C and 5% CO2. Cells were lysed, and mRNA quantification was performed using the QuantiGene 2.0 assay kit (Affymetrix).

### Lipid-mediated delivery

HeLa cells were plated in DMEM (Cellgro, #10-013CV) with 6% FBS (Gibco, #26140) at 10,000 cells per well in 96-well tissue culture-treated plates. SH-SY5Y cells (ATCC CRL-2266) were plated in EMEM (ATCC #30-2003) containing 6% FBS at 10,000 cells per well in 96-well tissue culture plates. Unconjugated siRNA was diluted in OptiMEM, and mixed 1:1 with Lipofectamine RNAiMAX Transfection Reagent (Invitrogen^TM^, #13778150). The final transfection reagent concentration was 0.3 μl/25 μl/well. siRNA and transfection reagent were added to cells resulting in 3% FBS final. Cells were incubated for 72 h at 37°C and 5% CO2.

### Mouse colonies

All mouse experiments were done following the regulations and guidelines set forth by the UMass Chan Medical School Institutional Animal Care and Use Committee, Protocol 202000010. Male and female mice, B6SJL-Tg(SOD1*G93A)1Gur/J (#002726), were obtained from Jackson Laboratory. They were allowed to acclimate to their new environment 1-2 weeks before injections.

### Stereotaxic intracerebroventricular injection

Mice were first anesthetized with Isoflurane (2.5% and 1.0L O2) in an induction chamber. Before being transferred to a nose cone, their heads were shaved and they were secured in the frame. A small incision was made exposing the skull, and the syringe zeroed over bregma. From there, coordinates were dialed for injection into the lateral ventricles. Injections were bilateral, with 5μl administered to each side. For repeat injection experiments, the initial injection was bilateral, with the 2 subsequent injections given unilaterally, one to each side. Post injection, the incision was sutured, analgesics given, and mice returned to a clean, warm cage where they are observed until fully awake, sternal, and mobile.

### Weight and Grip Strength

Mouse weight was recorded once per week. Grip-strength monitors the progression of clinical weakness in mouse models of neurodegeneration. Using a Mark-10 Digital Force Gauge, the strength of both front and hind limbs of mice were measured. The mice are manually held up to the wire mesh platform by their tail and allowed to grip it with their front paws. The mice are then gently pulled back by hand to measure their strength. Three trials are done with peak measurement taken. This is then repeated, allowing all four paws to grip the platform. Mice are tested once a week.

### ALS Motor Disease Score

Once a week, mice are given a score from 0-5 to evaluate disease progression. The basis for each score is as follows: 0: Healthy, clinically unremarkable. Full extension of hind legs away from lateral midline when suspended by its tail; 1: Collapse or partial collapse of leg extension towards lateral midline *and/or* weak hind limb tremor during tail suspension; 2: Definite tremor *and* lack of hind limb abduction; 3: Initial manifestation of paralysis displayed by decreased range of limb motion. Hind legs are bent and pulled inward. Gait may resemble a waddle; 4: Paralysis in one or more hind limbs. Legs may start to extend behind mouse. Foot may be delayed in flipping up when walking; 5: Mouse endpoint. Mouse cannot right itself within 10 seconds after being placed on its back.

### mRNA quantification

mRNA was quantified using the QuantiGene 2.0 Assay (Affymetrix, #QS0011). After 72-hour treatment, cells were lysed in 250 µL diluted lysis mixture with proteinase K (Affymetrix, #13228) for 30 min at 55°C. Probe sets and lysate amounts were validated to be in the linear range and were diluted as specified in the manufacturer’s protocol. Luminescence was detected on either a Veritas Luminometer (Promega, #998-9100) or a Tecan M1000 (Tecan, Morrisville, NC, USA). The specific mRNAs detected are specified in each graph. Catalog numbers are as follows: human HTT (Invitrogen^TM^, #SA-50339), human SOD1 (Invitrogen^TM^, #SA-10232), and human HPRT (Invitrogen^TM^, #SA-10030).

### qPCR analysis

RNA was extracted from cortex and cervical spinal cord tissue using Trizol Reagent (Life Technologies) following manufacturer’s protocols. 20 ng of purified RNA was added per well into a 384-well qPCR plate (Biorad) using Fast universal qPCR master mix (Applied Biosystems). Samples were amplified using a CFX384 Real Time System (Biorad). FAM labelled probes used during amplification included probes targeting Human SOD1 and Mouse HPRT (Applied Biosystems). Gene expression was calculated using the ΔΔCT method.

### Western Blot analysis

Protein from cortex and cervical spinal cord samples were extracted using RIPA buffer. 5µg of total protein was run per well in a 12-well 12% Tris-Glycine gel at 120 volts. Proteins were transferred to nitrocellulose paper using the iBlot 2 device (Invitrogen). Blots were incubated overnight in Intercept blocking buffer (Licor) containing a rabbit Anti-SOD1 antibody (GeneTex, GTX10054) and a goat anti-Anti β-Actin antibody (Abcam, ab8229). The next morning, blots were washed in PBS-T and incubated IR labelled infrared antibodies (donkey anti rabbit 680RD 926-32214, and donkey anti goat 800 CW 926-68073) for 1 hour at room temperature. After secondary incubation, blots were imaged using an Odyssey infrared imager (Licor).

### Immunohistochemistry

Brains and spinal cords were removed, postfixed overnight in 4% paraformaldehyde, and then stored in 0.4% paraformaldehyde at 4°C until further processing. Prior to paraffin embedding, brains were pre-sectioned using a brain matrix. For Nissl and immunohistochemistry assessment, paraffin brain sections, 10-µm thick sagittal, were obtained at approximately Bregma 1.08 mm and lumbosacral enlargement. Immunohistochemistry was performed against glial fibrillary acidic protein (GFAP, 1:250, Agilent, Cat# Z0334, RRID: AB_10013382), ionized calcium binding adaptor molecule 1 (Iba-1, 1:250, Wako, Cat# 019-19741, RRID: AB_839504), postsynaptic density protein 95 (PSD-95, 1:250, Thermo Fisher Scientific, Cat# MA1-045, RRID: AB_325399), and synaptophysin (Synaptophysin, 1:250, Millipore Cat# AB9272, RRID: AB_570874). For immunofluorescence staining, tissue sections labeled with the primary antibodies (GFAP, Iba-1, PSD-95, synaptophysin) were incubated in appropriate secondary antibodies conjugated with Alexa Fluor 488 (1:250, Abcam, Cat# ab150113, RRID: AB_2576208 and Cat# ab150077, RRID: AB_2630356), Alexa Fluor 555 (1:250, Abcam, Cat# ab150106, RRID: AB_2857373), and Alexa Fluor 647 (1:250, Abcam, Cat# ab150075, RRID: AB_2752244 and Cat# ab150115, RRID: AB_2687948). Omitting the primary antibody in a subset of slides served as negative controls. Mice at the following ages were used for histological analysis: naïve B6SJL, 160 days (14278: 170 days; 14270: 150 days); hSOD1^WT^, 145 days (14280: 145 days; 14284: 145 days); hSOD1^G93A^ (no tx), 87 days (14204: 100 days; 14192: 74 days); hSOD1^G93A^ (treated), 230 days (221: 230 days; 228: 230 days).

### Image acquisition and quantification

To acquire images of all stained sections for subsequent offline analysis, we used a Leica DM6 B microscopy system equipped with a brightfield DMC5400 color CMOS camera and an immunofluorescent DFC9000 sCMOS camera. For quantitative threshold area measurements of histological data, we used the Analyze Particle tool in ImageJ as described (*64*), with the experimenter blinded to the experimental groups (EOD). To determine the extent of neuronal loss, regions of interest (ROI: 329 x 219 µm) covering layer V of primary motor cortex (number of total neurons and dark neurons) and anterior horn of lumbar spinal cord (number of pyramidal neurons) were taken at 40x magnification using Nissl-stained images. To assess microgliosis and astrocytosis in layer V of primary motor cortex and anterior horn of lumbar spinal cord, we quantified the total thresholded area (ROI: 333 x 333 µm) of the Iba-1 and GFAP stained images (images taken at 40x magnification). For thresholded area (ROI: 211 x 211 µm) measurement of synaptophysin and PSD-95, images centered within the corresponding FOV used for the GFAP and Iba-1 analyses were taken at 63x magnification and analyzed as described for GFAP.

### Gastrocnemius neuromuscular junction

Mice assessed included untreated transgenic SOD1^WT^ and SOD1^G93A^, untreated non-transgenic B6SJL, and treated SOD1^G93A^, all of which are in the B6SJL background. Mice were transcardially perfused with PBS1X followed by 4% PFA. Gastrocnemius muscle extracted from fixed animal and placed in 5ml 4% PFA overnight then 5ml 30% sucrose overnight until sunk. Specimens embedded in OCT and sagittal sections were cut at 30 μm thickness and kept at -80°C until used. Antigen retrieval buffer (10mM sodium citrate dihydrate, 0.1% Tween-2, pH 6.5) was added to sections for 10 minutes at 95-100°C. Slides were then washed in PBS-T (0.1% tween in PBS1X) 3x for 5 minutes, and placed in blocking solution (10% normal donkey serum and 0.4% Triton in PBS1X) for 3 hours at room temperature. Placed in primary antibodies (Rabbit Beta-III Tubulin 1:500 (Covance, #PRB-435P-100), Rabbit Synaptophysin 1:500 (Invitrogen, #080130) diluted in blocking solution overnight-24 hours at 4°C then washed in PBS1X 3x for 5 minutes. Slides were then placed in secondary antibodies, Alpha-Bungarotoxin AF555 1:500 (ThermoFischer, #B35451) and Donkey anti-Rabbit AF488 1:450 (Jackson ImmunoResearch, #711-545-152), diluted in PBS1X for 2.5 hours at room temperature, then washed in PBS1X 3x for 5 minutes. Slides were mounted with Vectashield media with DAPI (Vector Laboratories, # H-1200-10). Neuromuscular junctions (NMJ) were blindly determined to be intact (90-100% colocalization), partially denervated (20-90% colocalization), or denervated (0-20% colocalization).

### Statistical analysis

Statistical analysis was performed with Graph Pad Prism Version 10.0.03. When comparing one variable, one-way ANOVA was used. When comparing two or more variables, two-way ANOVA with multiple comparisons was performed. Mantel-Cox test was performed to compare Kaplan-Meier curves. When reporting statistics, * p< 0.05, ** p< 0.01, *** p< 0.001, **** p < 0.0001.

## List of Supplementary Materials

Fig. S1: SOD1_123 silences both human and non-human primate mRNA *in vitro*.

Fig. S2: Chemical optimization of SOD1_123 reveals multiple potent configurations.

Fig. S3: Astrogliosis in the cortex but not lumbar spinal cord of di-siRNA treated G93A mice versus untreated G93A mice.

Fig. S4: Di-siRNA treatment activates cortical microglia and attenuates microgliosis in the anterior horn.

Table S1: siRNA sequences used for *in vitro* and *in vivo* studies

## ACKNOWLEDGMENTS

The authors thank Emily Haberlin for feedback on the manuscript.

## Funding

National Institutes of Health R35 GM131839 (AK)

National Institutes of Health R01 NS104022 (AK, RHB)

National Institutes of Health S10 OD020012 (AK).

National Institutes of Health NINDS R01 NS111990 (AK)

ALS Association (RHB)

The Angel Fund for ALS Research (RHB)

The Pierre L. de Bourgknecht ALS Research Fund (RHB)

ALS Finding A Cure (RHB)

ALSOne (RHB)

The Cellucci Fund (RHB)

The Max Rosenfeld ALS Research Fund (RHB)

The Ricci Fund for ALS Therapy Development (RHB)

National Institutes of Health NS131756 (NH)

National Institutes of Health NR020231 (NH)

National Institutes of Health U24NS113844 (NH).

Smith Family Foundation (PLG)

The Rita Allen Foundation (PLG)

Target ALS (PLG)

National Institutes of Health DP2 OD027719-01 (PLG)

## Author contributions

Conception and design: AK, RHB, BG, JG

Development of methodology: BG, MH, AW, JG, VR, CF, DE, BB, NM, NY, DC, KM, KY

Acquisition of data: AW, JG, VR, JB, CF, EOD, NW, BG, KM, AS, RF.

Analysis and interpretation of data: AW, JG, VR, JB, CF, EOD, NW, KM, NH, PLG

Writing – original draft: JB, RHB, NH, AK, VR, JG, AW, EOD, KM

Writing – review & editing: JB, NH, RHB, AK, VR, AW, JG, CF, EOD, NW, KM, AS, BB, NM, RF, NY, BG, MH, DC, FM, KY, PLG

Supervision of study: AK, RHB, NH, PLG

## Data and materials availability

Data is available upon request to the corresponding authors.

