## Supplemental Data for "RNAi-mediated silencing of SOD1 profoundly extends survival and functional outcomes in ALS mice"

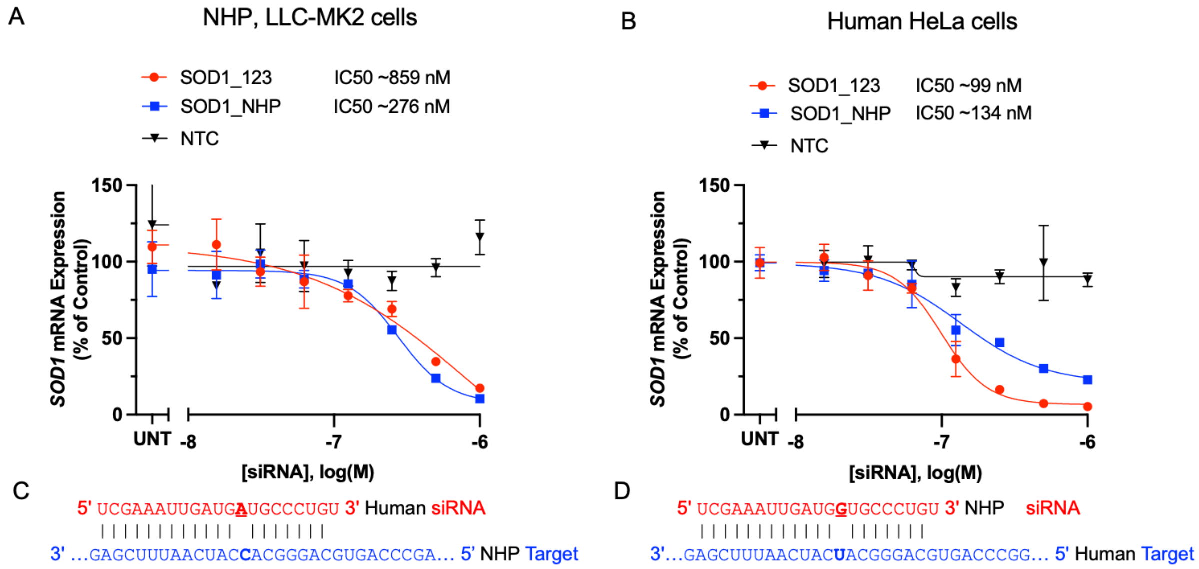

**Supplemental Figure 1: SOD1_123 silences both human and non-human primate mRNA *in vitro*.** (**A**, **B**) 7-point dose-response curves for SOD1_123 and SOD1_NHP in (**A**) LLC-MK2 cells (NHP-derived) or (**B**) HeLa cells (human-derived) (n = 3, mean ± SD). HeLa and LLC_MK2 cells were treated with siRNAs at concentrations shown for 72 h (Passive uptake). mRNA levels were measured using QuantiGene, (n=3, mean ± SD), UNT (untreated), NTC (non-targeting control). (**C**) Sequence of the human-targeting siRNA guide (red) against the NHP target mRNA (blue). Mismatch shown in bold. (**D**) Sequence of the NHP-targeting guide strand (red) against the human target mRNA (blue). Mismatch shown in bold. $IC_{50}$ values are shown above the graph. $IC_{50}$ values were calculated using the nonlinear least squares method (GraphPad Prism).

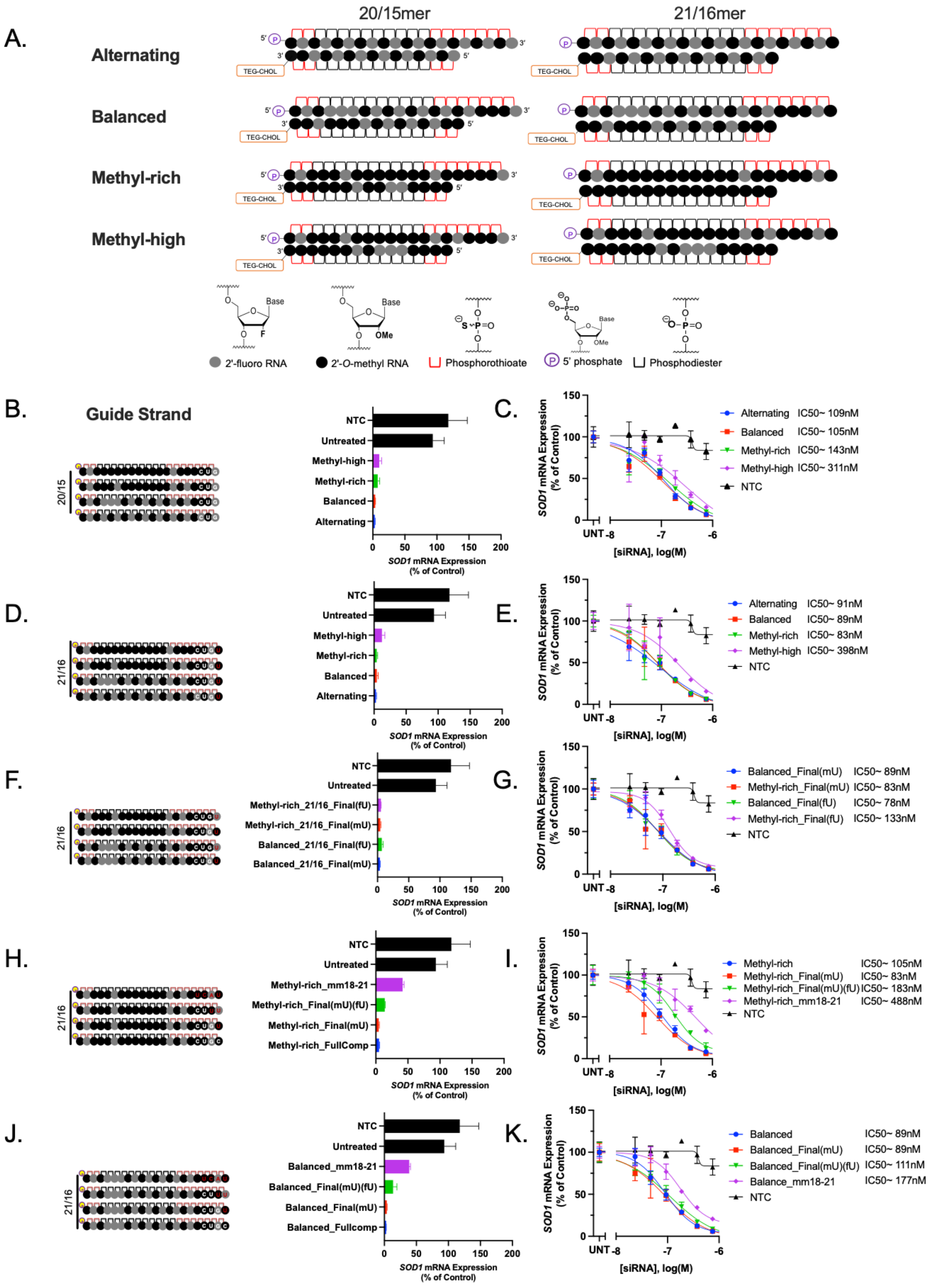

**Supplemental Figure 2: Chemical optimization of SOD1_123 reveals multiple potent configurations. (A)** Schematic of siRNA patterns (top), and chemical modifications (bottom). **(B)**

HeLa cells treated by passive uptake with SOD1_123 in four distinct patterns. The left panel shows diagram of the guide strands. **(C)** 7-point dose-reponse curves for siRNAs in B. **(D)** HeLa cells treated (by passive uptake) with SOD1_123 **(E)** 7-point dose-response curves for siRNAs described in D. **(F)** HeLa cells treated (by passive uptake) with the modification patterns shown **(G)** 7-point dose-response curves for siRNAs described in F. **(H)** HeLa cells treated (by passive uptake) with four SOD1_123 siRNAs in the Methyl-rich pattern, **(I)** 7-point dose-response curves for siRNAs described in F. **(J)** As in H but in the Balanced pattern. **(K)** Dose-dependent analysis of siRNAs described in H but in the Balanced pattern. Top dose 1.5 μM, SOD1 mRNA evaluated at 72 hours, QuantiGene, (n=3, mean ± SD). NTC (nontargeting control siRNA), UNT (untreated). The target site and $IC_{50}$ values are shown in the graphs. $IC_{50}$ values were calculated using the nonlinear least squares method (GraphPad Prism).

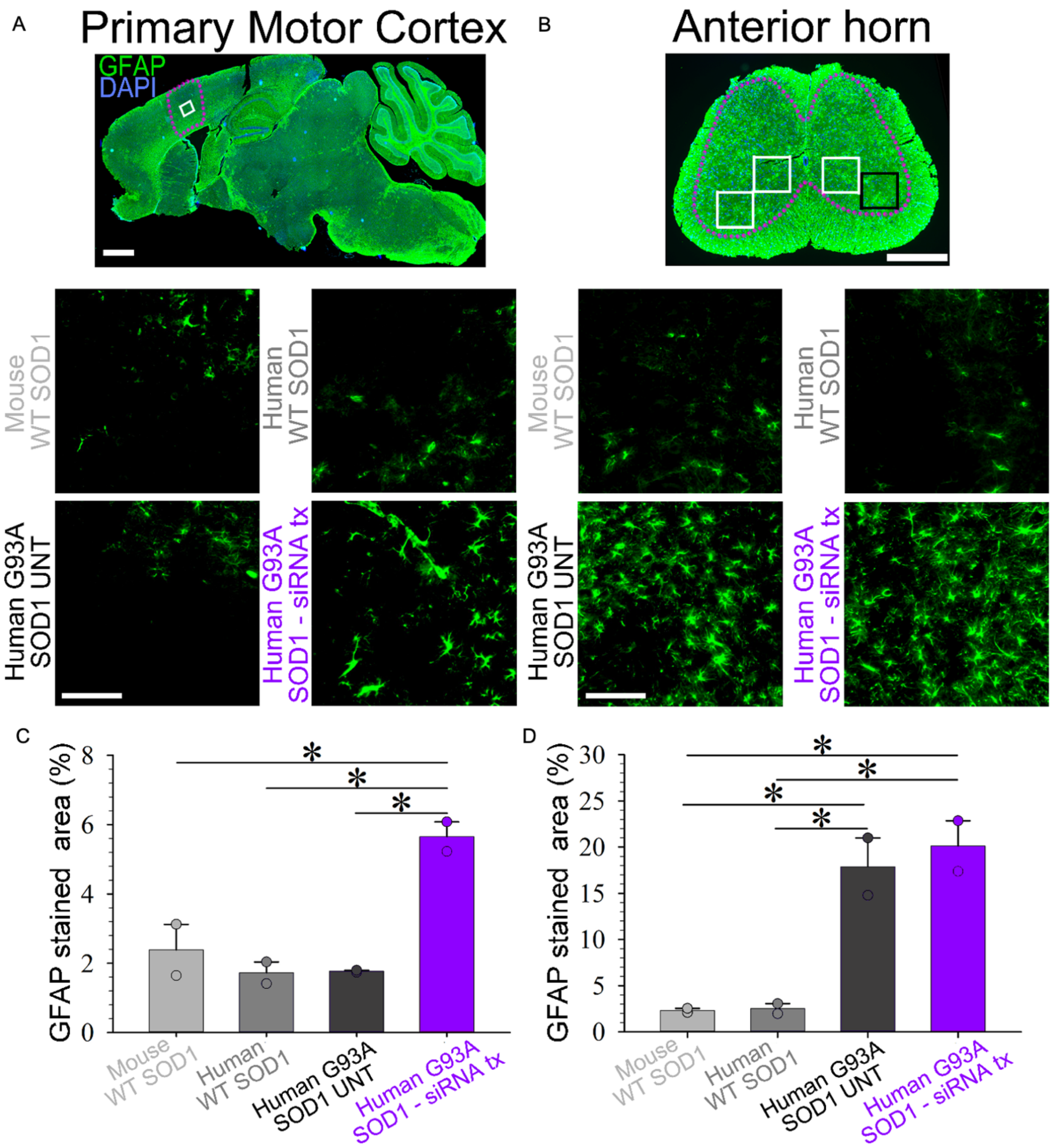

**Supplemental Figure 3: Astrogliosis in the cortex but not lumbar spinal cord of di-siRNA treated G93A mice versus untreated G93A mice. (A, B)** GFAP-stained sagittal section of brain (**A**) and transverse lumbar spinal cord (**B**) images with rectangles indicate the regions of interest (ROIs) used for quantification. **(C)** Representative images of layer V of primary motor cortex and anterior horn. Scale bar=100 μm. **(D, E)** Quantification of GFAP signal in layer V of primary motor cortex (**D**) and anterior horn of lumbar spinal cord (**E**). Data in the bar graphs are shown as mean ±SEM n=2 per group. One-way ANOVA with post hoc Holm-Šídák test.

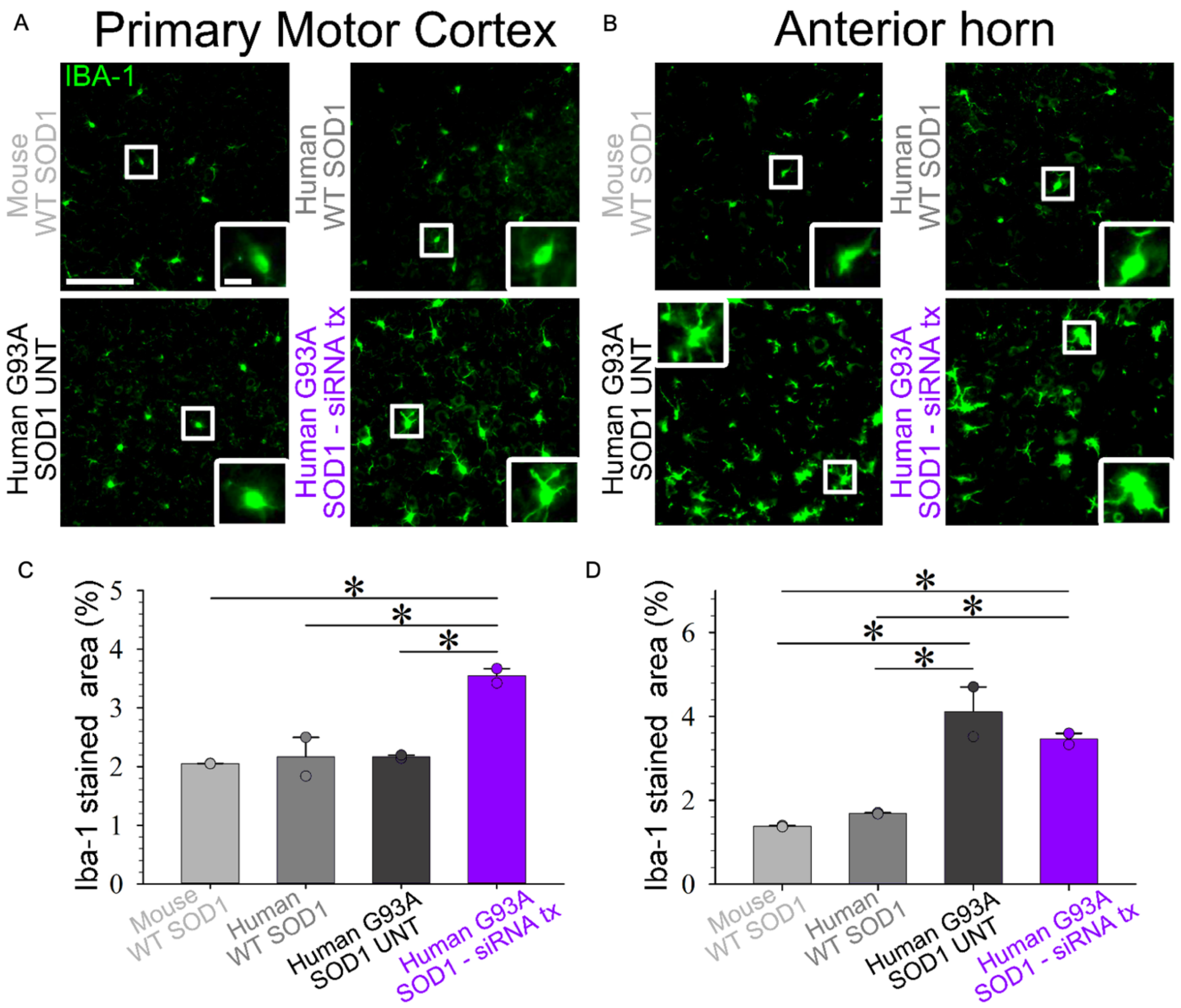

**Supplemental Figure 4: Di-siRNA treatment activates cortical microglia and attenuates microgliosis in the anterior horn. (A)** Representative images and insets of Iba-1 stained microglia in Motor cortex. Scale bar=100 µm Scale bar=15 µm (inset). **(B)** Representative images and insets of Iba-1 stained microglia in anterior horn. Scale bar=100 µm, scale bar in inset=15 µm. **(C)** Quantification of Iba-1 stained microglia in layer V of primary motor cortex and (**D**) anterior horn of lumbar spinal cord. Data in the bar graphs are shown as mean ±SEM n=2 per group. One-way ANOVA with post hoc Holm-Šídák test.

### **Supplemental Table 1:** siRNA sequences used for *in vitro* and *in vivo* studies

| Experiment | Antisense Strand (5'-3') | Sense Strand (5'-3') |
|---|---|---|
| Original SOD1 In Vitro Screen | | |
| siRNA Name | | |
| SOD1_796 | P(mU)#(fU)#(mA)(fA)(mC)(fU)(mG)(fA)(mG)(fU)(mU)(fU)(mU)#(fA)#(mU)#(fA)#(mA)#(fA)#(mA)#(fC) | (fA)#(mU)#(fA)(mA)(fA)(mA)(fC)(mU)(fC)(mA)(fG)(mU)(fU)#(mA)#(fA)-TegChol |
| SOD1_786 | P(mU)#(fU)#(mU)(fA)(mU)(fA)(mA)(fA)(mA)(fC)(mU)(fA)(mU)#(fA)#(mC)#(fA)#(mA)#(fA)#(mU)#(fC) | (fG)#(mU)#(fA)(mU)(fA)(mG)(fU)(mU)(fU)(mU)(fA)(mU)(fA)#(mA)#(fA)-TegChol |
| SOD1_595 | P(mU)#(fU)#(mU)(fA)(mU)(fU)(mG)(fG)(mG)(fC)(mG)(fA)(mU)#(fC)#(mC)#(fC)#(mA)#(fA)#(mU)#(fU) | (fG)#(mG)#(fA)(mU)(fC)(mG)(fC)(mC)(fC)(mA)(fA)(mU)(fA)#(mA)#(fA)-TegChol |
| SOD1_205 | P(mU)#(fU)#(mU)(fC)(mC)(fU)(mU)(fC)(mU)(fG)(mC)(fU)(mC)#(fG)#(mA)#(fA)#(mA)#(fU)#(mU)#(fG) | (fU)#(mC)#(fG)(mA)(fG)(mC)(fA)(mG)(fA)(mA)(fG)(mG)(fA)#(mA)#(fA)-TegChol |
| SOD1_926 | P(mU)#(fU)#(mU)(fA)(mA)(fU)(mA)(fG)(mC)(fC)(mU)(fC)(mA)#(fU)#(mA)#(fA)#(mU)#(fA)#(mA)#(fG) | (fU)#(mA)#(fU)(mG)(fA)(mG)(fG)(mC)(fU)(mA)(fU)(mU)(fA)#(mA)#(fA)-TegChol |
| SOD1_406 | P(mU)#(fU)#(mU)(fU)(mG)(fU)(mC)(fA)(mG)(fC)(mA)(fG)(mU)#(fC)#(mA)#(fC)#(mA)#(fU)#(mU)#(fG) | (fU)#(mG)#(fA)(mC)(fU)(mG)(fC)(mU)(fG)(mA)(fC)(mA)(fA)#(mA)#(fA)-TegChol |
| SOD1_754 | P(mU)#(fU)#(mA)(fA)(mA)(fU)(mC)(fA)(mG)(fU)(mU)(fU)(mC)#(fU)#(mC)#(fA)#(mC)#(fU)#(mA)#(fC) | (fG)#(mA)#(fG)(mA)(fA)(mA)(fC)(mU)(fG)(mA)(fU)(mU)(fU)#(mA)#(fA)-TegChol |
| SOD1_888 | P(mU)#(fU)#(mC)(fA)(mC)(fA)(mG)(fG)(mC)(fU)(mU)(fG)(mA)#(fA)#(mU)#(fG)#(mA)#(fC)#(mA)#(fA) | (fA)#(mU)#(fU)(mC)(fA)(mA)(fG)(mC)(fC)(mU)(fG)(mU)(fG)#(mA)#(fA)-TegChol |
| SOD1_915 | P(mU)#(fA)#(mU)(fA)(mA)(fU)(mA)(fA)(mG)(fU)(mG)(fC)(mC)#(fA)#(mU)#(fA)#(mC)#(fA)#(mG)#(fG) | (fA)#(mU)#(fG)(mG)(fC)(mA)(fC)(mU)(fU)(mA)(fU)(mU)(fA)#(mU)#(fA)-TegChol |
| SOD1_221 | P(mU)#(fU)#(mC)(fA)(mC)(fU)(mG)(fG)(mU)(fC)(mC)(fA)(mU)#(fU)#(mA)#(fC)#(mU)#(fU)#(mU)#(fC) | (fU)#(mA)#(fA)(mU)(fG)(mG)(fA)(mC)(fC)(mA)(fG)(mU)(fG)#(mA)#(fA)-TegChol |
| SOD1_879 | P(mU)#(fU)#(mG)(fA)(mA)(fU)(mG)(fA)(mC)(fA)(mA)(fA)(mG)#(fA)#(mA)#(fA)#(mU)#(fU)#(mC)#(fU) | (fU)#(mU)#(fC)(mU)(fU)(mU)(fG)(mU)(fC)(mA)(fU)(mU)(fC)#(mA)#(fA)-TegChol |
| SOD1_681 | P(mU)#(fG)#(mU)(fG)(mU)(fU)(mU)(fA)(mA)(fU)(mG)(fU)(mU)#(fU)#(mA)#(fU)#(mC)#(fA)#(mG)#(fG) | (fU)#(mA)#(fA)(mA)(fC)(mA)(fU)(mU)(fA)(mA)(fA)(mC)(fA)#(mC)#(fA)-TegChol |
| SOD1_744 | P(mU)#(fU)#(mC)(fU)(mC)(fA)(mC)(fU)(mA)(fC)(mA)(fG)(mG)#(fU)#(mA)#(fC)#(mU)#(fU)#(mU)#(fA) | (fU)#(mA)#(fC)(mC)(fU)(mG)(fU)(mA)(fG)(mU)(fG)(mA)(fG)#(mA)#(fA)-TegChol |
| SOD1_466 | P(mU)#(fG)#(mC)(fA)(mA)(fU)(mG)(fG)(mU)(fC)(mU)(fC)(mC)#(fU)#(mG)#(fA)#(mG)#(fA)#(mG)#(fU) | (fC)#(mA)#(fG)(mG)(fA)(mG)(fA)(mC)(fC)(mA)(fU)(mU)(fG)#(mC)#(fA)-TegChol |
| SOD1_893 | P(mU)#(fU)#(mU)(fU)(mA)(fU)(mU)(fC)(mA)(fC)(mA)(fG)(mG)#(fC)#(mU)#(fU)#(mG)#(fA)#(mA)#(fU) | (fA)#(mG)#(fC)(mC)(fU)(mG)(fU)(mG)(fA)(mA)(fU)(mA)(fA)#(mA)#(fA)-TegChol |
| SOD1_892 | P(mU)#(fU)#(mU)(fA)(mU)(fU)(mC)(fA)(mC)(fA)(mG)(fG)(mC)#(fU)#(mU)#(fG)#(mA)#(fA)#(mU)#(fG) | (fA)#(mA)#(fG)(mC)(fC)(mU)(fG)(mU)(fG)(mA)(fA)(mU)(fA)#(mA)#(fA)-TegChol |
| SOD1_821 | P(mU)#(fA)#(mU)(fA)(mC)(fA)(mG)(fG)(mU)(fC)(mA)(fU)(mU)#(fG)#(mA)#(fA)#(mA)#(fC)#(mA)#(fG) | (fU)#(mC)#(fA)(mA)(fU)(mG)(fA)(mC)(fC)(mU)(fG)(mU)(fA)#(mU)#(fA)-TegChol |
| SOD1_798 | P(mU)#(fU)#(mU)(fU)(mA)(fA)(mC)(fU)(mG)(fA)(mG)(fU)(mU)#(fU)#(mU)#(fA)#(mU)#(fA)#(mA)#(fA) | (fA)#(mA)#(fA)(mA)(fC)(mU)(fC)(mA)(fG)(mU)(fU)(mA)(fA)#(mA)#(fA)-TegChol |
| SOD1_657 | P(mU)#(fU)#(mU)(fU)(mC)(fU)(mA)(fC)(mA)(fG)(mC)(fU)(mA)#(fG)#(mC)#(fA)#(mG)#(fG)#(mA)#(fU) | (fG)#(mC)#(fU)(mA)(fG)(mC)(fU)(mG)(fU)(mA)(fG)(mA)(fA)#(mA)#(fA)-TegChol |
| SOD1_768 | P(mU)#(fC)#(mU)(fU)(mC)(fC)(mA)(fA)(mG)(fU)(mG)(fA)(mU)#(fC)#(mA)#(fU)#(mA)#(fA)#(mA)#(fU) | (fU)#(mG)#(fA)(mU)(fC)(mA)(fC)(mU)(fU)(mG)(fG)(mA)(fA)#(mG)#(fA)-TegChol |
| SOD1_750 | P(mU)#(fU)#(mC)(fA)(mG)(fU)(mU)(fU)(mC)(fU)(mC)(fA)(mC)#(fU)#(mA)#(fC)#(mA)#(fG)#(mG)#(fU) | (fU)#(mA)#(fG)(mU)(fG)(mA)(fG)(mA)(fA)(mA)(fC)(mU)(fG)#(mA)#(fA)-TegChol |
| SOD1_524 | P(mU)#(fU)#(mU)(fC)(mC)(fA)(mC)(fC)(mU)(fU)(mU)(fG)(mC)#(fC)#(mC)#(fA)#(mA)#(fG)#(mU)#(fC) | (fG)#(mG)#(fG)(mC)(fA)(mA)(fA)(mG)(fG)(mU)(fG)(mG)(fA)#(mA)#(fA)-TegChol |
| SOD1_252 | P(mU)#(fU)#(mC)(fA)(mG)(fU)(mC)(fA)(mG)(fU)(mC)(fC)(mU)#(fU)#(mU)#(fA)#(mA)#(fU)#(mG)#(fC) | (fA)#(mA)#(fA)(mG)(fG)(mA)(fC)(mU)(fG)(mA)(fC)(mU)(fG)#(mA)#(fA)-TegChol |
| SOD1_693 | P(mU)#(fU)#(mU)(fU)(mA)(fA)(mG)(fA)(mU)(fU)(mA)(fC)(mA)#(fG)#(mU)#(fG)#(mU)#(fU)#(mU)#(fA) | (fA)#(mC)#(fU)(mG)(fU)(mA)(fA)(mU)(fC)(mU)(fU)(mA)(fA)#(mA)#(fA)-TegChol |
| SOD1-535 | P(mU)#(fA)#(mC)(fU)(mU)(fU)(mC)(fU)(mU)(fC)(mA)(fU)(mU)#(fU)#(mC)#(fC)#(mA)#(fC)#(mC)#(fU) | (fG)#(mA)#(fA)(mA)(fU)(mG)(fA)(mA)(fG)(mA)(fA)(mA)(fG)#(mU)#(fA)-TegChol |
| SOD1_123 | P(mU)#(fC)#(mG)(mA)(mA)(fA)(mU)(mU)(mG)(mA)(mU)(mG)(mA)#(fU)#(mG)#(fC)#(mC)#(mC)#(mU)#(fG) | (mC)#(mA)#(mU)(mC)(fA)(fU)(fC)(mA)(fA)(mU)(mU)(mU)(mC)#(mG)#(mA)-TegChol |
| SOD1_249 | P(mU)#(fG)#(mA)(mG)(mG)(fA)(mC)(mC)(mU)(mG)(mC)(mA)(mC)#(fU)#(mG)#(fG)#(mU)#(mA)#(mC)#(fA) | (mC)#(mA)#(mG)(mU)(fG)(fC)(fA)(mG)(fG)(mU)(mC)(mC)(mU)#(mC)#(mA)-TegChol |
| SOD1_263 | P(mU)#(fU)#(mA)(mG)(mA)(fG)(mG)(mA)(mU)(mU)(mA)(mA)(mA)#(fG)#(mU)#(fG)#(mA)#(mG)#(mG)#(fA) | (mA)#(mC)#(mU)(mU)(fU)(fA)(fA)(mU)(fC)(mC)(mU)(mC)(mU)#(mA)#(mA)-TegChol |
| SOD1_345 | P(mU)#(fC)#(mC)(mA)(mC)(fA)(mC)(mC)(mA)(mU)(mC)(mU)(mU)#(fU)#(mG)#(fU)#(mC)#(mA)#(mG)#(fC) | (mC)#(mA)#(mA)(mA)(fG)(fA)(fU)(mG)(fG)(mU)(mG)(mU)(mG)#(mG)#(mA)-TegChol |
| SOD1_368 | P(mU)#(fA)#(mG)(mA)(mA)(fU)(mC)(mU)(mU)(mC)(mA)(mA)(mU)#(fA)#(mG)#(fA)#(mC)#(mA)#(mC)#(fA) | (mC)#(mU)#(mA)(mU)(fU)(fG)(fA)(mA)(fG)(mA)(mU)(mU)(mC)#(mU)#(mA)-TegChol |
| SOD1_384 | P(mU)#(fC)#(mU)(mG)(mA)(fG)(mA)(mG)(mU)(mG)(mA)(mG)(mA)#(fU)#(mC)#(fA)#(mC)#(mA)#(mG)#(fA) | (mG)#(mA)#(mU)(mC)(fU)(fC)(fA)(mC)(fU)(mC)(mU)(mC)(mA)#(mG)#(mA)-TegChol |
| SOD1_457 | P(mU)#(fU)#(mC)(mA)(mU)(fU)(mU)(mC)(mC)(mA)(mC)(mC)(mU)#(fU)#(mU)#(fG)#(mC)#(mC)#(mC)#(fA) | (mA)#(mA)#(mA)(mG)(fG)(fU)(fG)(mG)(fA)(mA)(mA)(mU)(mG)#(mA)#(mA)-TegChol |
| SOD1_516 | P(mU)#(fC)#(mG)(mA)(mU)(fC)(mC)(mC)(mA)(mA)(mU)(mU)(mA)#(fC)#(mA)#(fC)#(mC)#(mA)#(mC)#(fA) | (mU)#(mG)#(mU)(mA)(fA)(fU)(fU)(mG)(fG)(mG)(mA)(mU)(mC)#(mG)#(mA)-TegChol |
| SOD1_573 | P(mU)#(fC)#(mA)(mG)(mG)(fA)(mU)(mA)(mA)(mC)(mA)(mG)(mA)#(fU)#(mG)#(fA)#(mG)#(mU)#(mU)#(fA) | (mC)#(mA)#(mU)(mC)(fU)(fG)(fU)(mU)(fA)(mU)(mC)(mC)(mU)#(mG)#(mA)-TegChol |
| SOD1_594 | P(mU)#(fA)#(mG)(mG)(mA)(fU)(mA)(mC)(mA)(mU)(mU)(mU)(mC)#(fU)#(mA)#(fC)#(mA)#(mG)#(mC)#(fU) | (mU)#(mA)#(mG)(mA)(fA)(fA)(fU)(mG)(fU)(mA)(mU)(mC)(mC)#(mU)#(mA)-TegChol |
| SOD1_614 | P(mU)#(fU)#(mA)(mC)(mA)(fG)(mU)(mG)(mU)(mU)(mU)(mA)(mA)#(fU)#(mG)#(fU)#(mU)#(mU)#(mA)#(fU) | (mC)#(mA)#(mU)(mU)(fA)(fA)(fA)(mC)(fA)(mC)(mU)(mG)(mU)#(mA)#(mA)-TegChol |
| SOD1_627 | P(mU)#(fA)#(mC)(mA)(mC)(fU)(mU)(mU)(mU)(mA)(mA)(mG)(mA)#(fU)#(mU)#(fA)#(mC)#(mA)#(mG)#(fU) | (mA)#(mA)#(mU)(mC)(fU)(fU)(fA)(mA)(fA)(mA)(mG)(mU)(mG)#(mU)#(mA)-TegChol |
| SOD1_636 | P(mU)#(fC)#(mA)(mC)(mA)(fC)(mA)(mA)(mU)(mU)(mA)(mC)(mA)#(fC)#(mU)#(fU)#(mU)#(mU)#(mA)#(fA) | (mA)#(mG)#(mU)(mG)(fU)(fA)(fA)(mU)(fU)(mG)(mU)(mG)(mU)#(mG)#(mA)-TegChol |
| SOD1_681 | P(mU)#(fA)#(mA)(mU)(mC)(fA)(mG)(mU)(mU)(mU)(mC)(mU)(mC)#(fA)#(mC)#(fU)#(mA)#(mC)#(mA)#(fG) | (mG)#(mU)#(mG)(mA)(fG)(fA)(fA)(mA)(fC)(mU)(mG)(mA)(mU)#(mU)#(mA)-TegChol |
| SOD1_682 | P(mU)#(fA)#(mA)(mA)(mU)(fC)(mA)(mG)(mU)(mU)(mU)(mC)(mU)#(fC)#(mA)#(fC)#(mU)#(mA)#(mC)#(fA) | (mU)#(mG)#(mA)(mG)(fA)(fA)(fA)(mC)(fU)(mG)(mA)(mU)(mU)#(mU)#(mA)-TegChol |
| SOD1_701 | P(mU)#(fA)#(mA)(mA)(mU)(fC)(mU)(mU)(mC)(mC)(mA)(mA)(mG)#(fU)#(mG)#(fA)#(mU)#(mC)#(mA)#(fU) | (mC)#(mA)#(mC)(mU)(fU)(fG)(fG)(mA)(fA)(mG)(mA)(mU)(mU)#(mU)#(mA)-TegChol |
| SOD1_710 | P(mU)#(fA)#(mA)(mA)(mC)(fU)(mA)(mU)(mA)(mC)(mA)(mA)(mA)#(fU)#(mC)#(fU)#(mU)#(mC)#(mC)#(fA) | (mG)#(mA)#(mU)(mU)(fU)(fG)(fU)(mA)(fU)(mA)(mG)(mU)(mU)#(mU)#(mA)-TegChol |
| SOD1_724 | P(mU)#(fA)#(mA)(mC)(mU)(fG)(mA)(mG)(mU)(mU)(mU)(mU)(mA)#(fU)#(mA)#(fA)#(mA)#(mA)#(mC)#(fU) | (mU)#(mA)#(mU)(mA)(fA)(fA)(fA)(mC)(fU)(mC)(mA)(mG)(mU)#(mU)#(mA)-TegChol |
| SOD1_752 | P(mU)#(fA)#(mA)(mA)(mU)(fA)(mC)(mA)(mG)(mG)(mU)(mC)(mA)#(fU)#(mU)#(fG)#(mA)#(mA)#(mA)#(fC) | (mA)#(mA)#(mU)(mG)(fA)(fC)(fC)(mU)(fG)(mU)(mA)(mU)(mU)#(mU)#(mA)-TegChol |
| SOD1_769 | P(mU)#(fG)#(mU)(mG)(mA)(fU)(mU)(mU)(mA)(mA)(mG)(mU)(mC)#(fU)#(mG)#(fG)#(mC)#(mA)#(mA)#(fA) | (mC)#(mA)#(mG)(mA)(fC)(fU)(fU)(mA)(fA)(mA)(mU)(mC)(mA)#(mC)#(mA)-TegChol |
| SOD1_790 | P(mU)#(fU)#(mG)(mA)(mC)(fA)(mA)(mG)(mU)(mU)(mU)(mA)(mA)#(fU)#(mA)#(fC)#(mC)#(mC)#(mA)#(fU) | (mU)#(mA)#(mU)(mU)(fA)(fA)(fA)(mC)(fU)(mU)(mG)(mU)(mC)#(mA)#(mA)-TegChol |
| SOD1_797 | P(mU)#(fG)#(mA)(mA)(mA)(fU)(mU)(mC)(mU)(mG)(mA)(mC)(mA)#(fA)#(mG)#(fU)#(mU)#(mU)#(mA)#(fA) | (mC)#(mU)#(mU)(mG)(fU)(fC)(fA)(mG)(fA)(mA)(mU)(mU)(mU)#(mC)#(mA)-TegChol |
| SOD1_813 | P(mU)#(fA)#(mG)(mG)(mC)(fU)(mU)(mG)(mA)(mA)(mU)(mG)(mA)#(fC)#(mA)#(fA)#(mA)#(mG)#(mA)#(fA) | (mU)#(mG)#(mU)(mC)(fA)(fU)(fU)(mC)(fA)(mA)(mG)(mC)(mC)#(mU)#(mA)-TegChol |
| SOD1_848 | P(mU)#(fC)#(mC)(mU)(mC)(fA)(mU)(mA)(mA)(mU)(mA)(mA)(mG)#(fU)#(mG)#(fC)#(mC)#(mA)#(mU)#(fA) | (mC)#(mA)#(mC)(mU)(fU)(fA)(fU)(mU)(fA)(mU)(mG)(mA)(mG)#(mG)#(mA)-TegChol |
| SOD1_123 Walk | | |
| SOD1_112 | P(mU)#(fA)#(mU)(fG)(fC)(fC)(mC)(fU)(mG)(fC)(mA)(fC)(mU)(fG)#(mG)#(fG)#(mC)#(mC)#(mG)#(fU)#(mU) | (mC)#(mC)#(mC)(fA)(mG)(fU)(mG)(fC)(mA)(fG)(mG)(mG)(mC)(fA)#(mU)#(mA)-TegChol |
| SOD1_113 | P(mU)#(fG)#(mA)(fU)(fG)(fC)(mC)(fC)(mU)(fG)(mC)(fA)(mC)(fU)#(mG)#(fG)#(mG)#(mC)#(mC)#(fG)#(mU) | (mC)#(mC)#(mA)(fG)(mU)(fG)(mC)(fA)(mG)(fG)(mG)(mC)(mA)(fU)#(mC)#(mA)-TegChol |
| SOD1_114 | P(mU)#(fU)#(mG)(fA)(fU)(fG)(mC)(fC)(mC)(fU)(mG)(fC)(mA)(fC)#(mU)#(fG)#(mG)#(mG)#(mC)#(fC)#(mU) | (mC)#(mA)#(mG)(fU)(mG)(fC)(mA)(fG)(mG)(fG)(mC)(mA)(mU)(fC)#(mA)#(mA)-TegChol |
| SOD1_115 | P(mU)#(fA)#(mU)(fG)(fA)(fU)(mG)(fC)(mC)(fC)(mU)(fG)(mC)(fA)#(mC)#(fU)#(mG)#(mG)#(mG)#(fC)#(mU) | (mA)#(mG)#(mU)(fG)(mC)(fA)(mG)(fG)(mG)(fC)(mA)(mU)(mC)(fA)#(mU)#(mA)-TegChol |
| SOD1_116 | P(mU)#(fG)#(mA)(fU)(fG)(fA)(mU)(fG)(mC)(fC)(mC)(fU)(mG)(fC)#(mA)#(fC)#(mU)#(mG)#(mG)#(fG)#(mU) | (mG)#(mU)#(mG)(fC)(mA)(fG)(mG)(fG)(mC)(fA)(mU)(mC)(mA)(fU)#(mC)#(mA)-TegChol |
| SOD1_117 | P(mU)#(fU)#(mG)(fA)(fU)(fG)(mA)(fU)(mG)(fC)(mC)(fC)(mU)(fG)#(mC)#(fA)#(mC)#(mU)#(mG)#(fG)#(mU) | (mU)#(mG)#(mC)(fA)(mG)(fG)(mG)(fC)(mA)(fU)(mC)(mA)(mU)(fC)#(mA)#(mA)-TegChol |
| SOD1_118 | P(mU)#(fU)#(mU)(fG)(fA)(fU)(mG)(fA)(mU)(fG)(mC)(fC)(mC)(fU)#(mG)#(fC)#(mA)#(mC)#(mU)#(fG)#(mU) | (mG)#(mC)#(mA)(fG)(mG)(fG)(mC)(fA)(mU)(fC)(mA)(mU)(mC)(fA)#(mA)#(mA)-TegChol |
| SOD1_119 | P(mU)#(fA)#(mU)(fU)(fG)(fA)(mU)(fG)(mA)(fU)(mG)(fC)(mC)(fC)#(mU)#(fG)#(mC)#(mA)#(mC)#(fU)#(mU) | (mC)#(mA)#(mG)(fG)(mG)(fC)(mA)(fU)(mC)(fA)(mU)(mC)(mA)(fA)#(mU)#(mA)-TegChol |
| SOD1_120 | P(mU)#(fA)#(mA)(fU)(fU)(fG)(mA)(fU)(mG)(fA)(mU)(fG)(mC)(fC)#(mC)#(fU)#(mG)#(mC)#(mA)#(fC)#(mU) | (mA)#(mG)#(mG)(fG)(mC)(fA)(mU)(fC)(mA)(fU)(mC)(mA)(mA)(fU)#(mU)#(mA)-TegChol |
| SOD1_121 | P(mU)#(fA)#(mA)(fA)(fU)(fU)(mG)(fA)(mU)(fG)(mA)(fU)(mG)(fC)#(mC)#(fC)#(mU)#(mG)#(mC)#(fA)#(mU) | (mG)#(mG)#(mG)(fC)(mA)(fU)(mC)(fA)(mU)(fC)(mA)(mA)(mU)(fU)#(mU)#(mA)-TegChol |
| SOD1_122 | P(mU)#(fG)#(mA)(fA)(fA)(fU)(mU)(fG)(mA)(fU)(mG)(fA)(mU)(fG)#(mC)#(fC)#(mC)#(mU)#(mG)#(fC)#(mU) | (mG)#(mG)#(mC)(fA)(mU)(fC)(mA)(fU)(mC)(fA)(mA)(mU)(mU)(fU)#(mC)#(mA)-TegChol |
| SOD1_123 | P(mU)#(fC)#(mG)(fA)(fA)(fA)(mU)(fU)(mG)(fA)(mU)(fG)(mA)(fU)#(mG)#(fC)#(mC)#(mC)#(mU)#(fG)#(mU) | (mG)#(mC)#(mA)(fU)(mC)(fA)(mU)(fC)(mA)(fA)(mU)(mU)(mU)(fC)#(mG)#(mA)-TegChol |
| SOD1_124 | P(mU)#(fU)#(mC)(fG)(fA)(fA)(mA)(fU)(mU)(fG)(mA)(fU)(mG)(fA)#(mU)#(fG)#(mC)#(mC)#(mC)#(fU)#(mU) | (mC)#(mA)#(mU)(fC)(mA)(fU)(mC)(fA)(mA)(fU)(mU)(mU)(mC)(fG)#(mA)#(mA)-TegChol |
| SOD1_125 | P(mU)#(fC)#(mU)(fC)(fG)(fA)(mA)(fA)(mU)(fU)(mG)(fA)(mU)(fG)#(mA)#(fU)#(mG)#(mC)#(mC)#(fC)#(mU) | (mA)#(mU)#(mC)(fA)(mU)(fC)(mA)(fA)(mU)(fU)(mU)(mC)(mG)(fA)#(mG)#(mA)-TegChol |
| SOD1_126 | P(mU)#(fG)#(mC)(fU)(fC)(fG)(mA)(fA)(mA)(fU)(mU)(fG)(mA)(fU)#(mG)#(fA)#(mU)#(mG)#(mC)#(fC)#(mU) | (mU)#(mC)#(mA)(fU)(mC)(fA)(mA)(fU)(mU)(fU)(mC)(mG)(mA)(fG)#(mC)#(mA)-TegChol |
| SOD1_127 | P(mU)#(fU)#(mG)(fC)(fU)(fC)(mG)(fA)(mA)(fA)(mU)(fU)(mG)(fA)#(mU)#(fG)#(mA)#(mU)#(mG)#(fC)#(mU) | (mC)#(mA)#(mU)(fC)(mA)(fA)(mU)(fU)(mU)(fC)(mG)(mA)(mG)(fC)#(mA)#(mA)-TegChol |
| SOD1_128 | P(mU)#(fC)#(mU)(fG)(fC)(fU)(mC)(fG)(mA)(fA)(mA)(fU)(mU)(fG)#(mA)#(fU)#(mG)#(mA)#(mU)#(fG)#(mU) | (mA)#(mU)#(mC)(fA)(mA)(fU)(mU)(fU)(mC)(fG)(mA)(mG)(mC)(fA)#(mG)#(mA)-TegChol |
| SOD1_129 | P(mU)#(fU)#(mC)(fU)(fG)(fC)(mU)(fC)(mG)(fA)(mA)(fA)(mU)(fU)#(mG)#(fA)#(mU)#(mG)#(mA)#(fU)#(mU) | (mU)#(mC)#(mA)(fA)(mU)(fU)(mU)(fC)(mG)(fA)(mG)(mC)(mA)(fG)#(mA)#(mA)-TegChol |
| SOD1_130 | P(mU)#(fU)#(mU)(fC)(fU)(fG)(mC)(fU)(mC)(fG)(mA)(fA)(mA)(fU)#(mU)#(fG)#(mA)#(mU)#(mG)#(fA)#(mU) | (mC)#(mA)#(mA)(fU)(mU)(fU)(mC)(fG)(mA)(fG)(mC)(mA)(mG)(fA)#(mA)#(mA)-TegChol |
| SOD1_131 | P(mU)#(fC)#(mU)(fU)(fC)(fU)(mG)(fC)(mU)(fC)(mG)(fA)(mA)(fA)#(mU)#(fU)#(mG)#(mA)#(mU)#(fG)#(mU) | (mA)#(mA)#(mU)(fU)(mU)(fC)(mG)(fA)(mG)(fC)(mA)(mG)(mA)(fA)#(mG)#(mA)-TegChol |
| SOD1_132 | P(mU)#(fC)#(mC)(fU)(fU)(fC)(mU)(fG)(mC)(fU)(mC)(fG)(mA)(fA)#(mA)#(fU)#(mU)#(mG)#(mA)#(fU)#(mU) | (mA)#(mU)#(mU)(fU)(mC)(fG)(mA)(fG)(mC)(fA)(mG)(mA)(mA)(fG)#(mG)#(mA)-TegChol |
| SOD1_133 | P(mU)#(fU)#(mC)(fC)(fU)(fU)(mC)(fU)(mG)(fC)(mU)(fC)(mG)(fA)#(mA)#(fA)#(mU)#(mU)#(mG)#(fA)#(mU) | (mU)#(mU)#(mU)(fC)(mG)(fA)(mG)(fC)(mA)(fG)(mA)(mA)(mG)(fG)#(mA)#(mA)-TegChol |
| SOD1_134 | P(mU)#(fU)#(mU)(fC)(fC)(fU)(mU)(fC)(mU)(fG)(mC)(fU)(mC)(fG)#(mA)#(fA)#(mA)#(mU)#(mU)#(fG)#(mU) | (mU)#(mU)#(mC)(fG)(mA)(fG)(mC)(fA)(mG)(fA)(mA)(mG)(mG)(fA)#(mA)#(mA)-TegChol |
| SOD1_135 | P(mU)#(fU)#(mU)(fU)(fC)(fC)(mU)(fU)(mC)(fU)(mG)(fC)(mU)(fC)#(mG)#(fA)#(mA)#(mA)#(mU)#(fU)#(mU) | (mU)#(mC)#(mG)(fA)(mG)(fC)(mA)(fG)(mA)(fA)(mG)(mG)(mA)(fA)#(mA)#(mA)-TegChol |
| Tofesen comparison | | |
| Tofersen | (eC)#(eA)(eG)#(eG)(eA)#(dT)#(dA)#(dC)#(dA)#(dT)#(dT)#(dT)#(dC)#(dT)#(dA)#(eC)(eA)#(eG)(eC)#(eT) | |
| Non-Human Primate (NHP) Targeting | | |
| SOD1_NHP (position 47) | P(mU)#(fC)#(mG)(mA)(mA)(fA)(mU)(mU)(mG)(mA)(mU)(mG)(mG)(fU)#(mG)#(fC)#(mC)#(mC)#(mU)#(fG)#(mU) | (mG)#(mC)#(mA)(mC)(mC)(fA)(fU)(fC)(mA)(fA)(mU)(mU)(mU)(mC)#(mG)#(mA)-TegChol |
| Human SOD1_123 | P(mU)#(fC)#(mG)(mA)(mA)(fA)(mU)(mU)(mG)(mA)(mU)(mG)(mA)(fU)#(mG)#(fC)#(mC)#(mC)#(mU)#(fG)#(mU) | (mG)#(mC)#(mA)(mU)(mC)(fA)(fU)(fC)(mA)(fA)(mU)(mU)(mU)(mC)#(mG)#(mA)-TegChol |
| Chemical Optimization Experiments | | |
| Alternating 20/15 | P(mU)#(fC)#(mG)(fA)(mA)(fA)(mU)(fU)(mG)(fA)(mU)(fG)(mA)#(fU)#(mG)#(fC)#(mC)#(fC)#(mU)#(fG) | (fC)#(mA)#(fU)(mC)(fA)(mU)(fC)(mA)(fA)(mU)(fU)(mU)(fC)#(mG)#(mA)-TegChol |
| Balanced 20/15 | P(mU)#(fC)#(mG)(fA)(fA)(fA)(mU)(fU)(mG)(fA)(mU)(fG)(mA)#(fU)#(mG)#(fC)#(mC)#(mC)#(mU)#(fG) | (mC)#(mA)#(fU)(mC)(fA)(mU)(fC)(mA)(fA)(mU)(mU)(mU)(fC)#(mG)#(mA)-TegChol |
| Methyl-high 20/15 | P(mU)#(fC)#(mG)(mA)(mA)(mA)(mU)(mU)(mG)(mA)(mU)(mG)(mA)#(fU)#(mG)#(mC)#(mC)#(mC)#(mU)#(fG) | (mC)#(mA)#(mU)(mC)(mA)(mU)(mC)(mA)(mA)(mU)(mU)(mU)(mC)#(mG)#(mA)-TegChol |
| Methyl-rich 20/15 | P(mU)#(fC)#(mG)(mA)(mA)(fA)(mU)(mU)(mG)(mA)(mU)(mG)(mA)#(fU)#(mG)#(fC)#(mC)#(mC)#(mU)#(fG) | (mC)#(mA)#(mU)(mC)(fA)(fU)(fC)(mA)(fA)(mU)(mU)(mU)(mC)#(mG)#(mA)-TegChol |
| Alternating 21/16 Final(mU) | P(mU)#(fC)#(mG)(fA)(mA)(fA)(mU)(fU)(mG)(fA)(mU)(fG)(mA)(fU)#(mG)#(fC)#(mC)#(fC)#(mU)#(fG)#(mU) | (mG)#(fC)#(mA)(fU)(mC)(fA)(mU)(fC)(mA)(fA)(mU)(fU)(mU)(fC)#(mG)#(mA)-TegChol |
| Balanced 21/16 Final (mU) | P(mU)#(fC)#(mG)(fA)(fA)(fA)(mU)(fU)(mG)(fA)(mU)(fG)(mA)(fU)#(mG)#(fC)#(mC)#(mC)#(mU)#(fG)#(mU) | (mG)#(mC)#(mA)(fU)(mC)(fA)(mU)(fC)(mA)(fA)(mU)(mU)(mU)(fC)#(mG)#(mA)-TegChol |
| Methyl-high 21/16 Final (mU) | P(mU)#(fC)#(mG)(mA)(mA)(mA)(mU)(mU)(mG)(mA)(mU)(mG)(mA)(fU)#(mG)#(mC)#(mC)#(mC)#(mU)#(fG)#(mU) | (mG)#(mC)#(mA)(mU)(mC)(mA)(mU)(mC)(mA)(mA)(mU)(mU)(mU)(mC)#(mG)#(mA)-TegChol |
| Methyl-rich 21/16 Final (mU) | P(mU)#(fC)#(mG)(mA)(mA)(fA)(mU)(mU)(mG)(mA)(mU)(mG)(mA)(fU)#(mG)#(fC)#(mC)#(mC)#(mU)#(fG)#(mU) | (mG)#(mC)#(mA)(mU)(mC)(fA)(fU)(fC)(mA)(fA)(mU)(mU)(mU)(mC)#(mG)#(mA)-TegChol |
| Balanced 21/16 FullComp | P(mU)#(fC)#(mG)(fA)(fA)(fA)(mU)(fU)(mG)(fA)(mU)(fG)(mA)(fU)#(mG)#(fC)#(mC)#(mC)#(mU)#(fG)#(mC) | (mG)#(mC)#(mA)(fU)(mC)(fA)(mU)(fC)(mA)(fA)(mU)(mU)(mU)(fC)#(mG)#(mA)-TegChol |
| Methyl-rich 21/16 FullComp | P(mU)#(fC)#(mG)(mA)(mA)(fA)(mU)(mU)(mG)(mA)(mU)(mG)(mA)(fU)#(mG)#(fC)#(mC)#(mC)#(mU)#(fG)#(mC) | (mG)#(mC)#(mA)(mU)(mC)(fA)(fU)(fC)(mA)(fA)(mU)(mU)(mU)(mC)#(mG)#(mA)-TegChol |
| Balanced 20/15 mm18-20 | P(mU)#(fC)#(mG)(fA)(fA)(fA)(mU)(fU)(mG)(fA)(mU)(fG)(mA)#(fU)#(mG)#(fC)#(mC)#(mU)#(mC)#(fA) | (mC)#(mA)#(fU)(mC)(fA)(mU)(fC)(mA)(fA)(mU)(mU)(mU)(fC)#(mG)#(mA)-TegChol |
| Methyl-rich 20/15 mm18-20 | P(mU)#(fC)#(mG)(mA)(mA)(fA)(mU)(mU)(mG)(mA)(mU)(mG)(mA)#(fU)#(mG)#(fC)#(mC)#(mU)#(mC)#(fA) | (mC)#(mA)#(mU)(mC)(fA)(fU)(fC)(mA)(fA)(mU)(mU)(mU)(mC)#(mG)#(mA)-TegChol |
| Balanced 21/16 mm18-21 | P(mU)#(fC)#(mG)(fA)(fA)(fA)(mU)(fU)(mG)(fA)(mU)(fG)(mA)(fU)#(mG)#(fC)#(mC)#(mU)#(mC)#(fA)#(mU) | (mG)#(mC)#(mA)(fU)(mC)(fA)(mU)(fC)(mA)(fA)(mU)(mU)(mU)(fC)#(mG)#(mA)-TegChol |
| Methyl-rich 21/16 mm18-21 | P(mU)#(fC)#(mG)(mA)(mA)(fA)(mU)(mU)(mG)(mA)(mU)(mG)(mA)(fU)#(mG)#(fC)#(mC)#(mU)#(mC)#(fA)#(mU) | (mG)#(mC)#(mA)(mU)(mC)(fA)(fU)(fC)(mA)(fA)(mU)(mU)(mU)(mC)#(mG)#(mA)-TegChol |
| Methyl-rich 21/16 Final (mU) | P(mU)#(fC)#(mG)(mA)(mA)(fA)(mU)(mU)(mG)(mA)(mU)(mG)(mA)(fU)#(mG)#(fC)#(mC)#(mC)#(mU)#(fG)#(mU) | (mG)#(mC)#(mA)(mU)(mC)(fA)(fU)(fC)(mA)(fA)(mU)(mU)(mU)(mC)#(mG)#(mA)-TegChol |
| Methyl-rich 21/16 final (fU) | P(mU)#(fC)#(mG)(mA)(mA)(fA)(mU)(mU)(mG)(mA)(mU)(mG)(mA)(fU)#(mG)#(fC)#(mC)#(mC)#(mU)#(mG)#(fU) | (mG)#(mC)#(mA)(mU)(mC)(fA)(fU)(fC)(mA)(fA)(mU)(mU)(mU)(mC)#(mG)#(mA)-TegChol |
| Methyl-rich 21/16 mm20-21 | P(mU)#(fC)#(mG)(mA)(mA)(fA)(mU)(mU)(mG)(mA)(mU)(mG)(mA)(fU)#(mG)#(fC)#(mC)#(mC)#(mU)#(mU)#(fU) | (mG)#(mC)#(mA)(mU)(mC)(fA)(fU)(fC)(mA)(fA)(mU)(mU)(mU)(mC)#(mG)#(mA)-TegChol |
| Balanced 21/16 Final (mU) | P(mU)#(fC)#(mG)(fA)(fA)(fA)(mU)(fU)(mG)(fA)(mU)(fG)(mA)(fU)#(mG)#(fC)#(mC)#(mC)#(mU)#(fG)#(mU) | (mG)#(mC)#(mA)(fU)(mC)(fA)(mU)(fC)(mA)(fA)(mU)(mU)(mU)(fC)#(mG)#(mA)-TegChol |
| Balanced 21/16 Final (fU) | P(mU)#(fC)#(mG)(fA)(fA)(fA)(mU)(fU)(mG)(fA)(mU)(fG)(mA)(fU)#(mG)#(fC)#(mC)#(mC)#(mU)#(mG)#(fU) | (mG)#(mC)#(mA)(fU)(mC)(fA)(mU)(fC)(mA)(fA)(mU)(mU)(mU)(fC)#(mG)#(mA)-TegChol |
| Balanced 21/16 mm20-21 | P(mU)#(fC)#(mG)(fA)(fA)(fA)(mU)(fU)(mG)(fA)(mU)(fG)(mA)(fU)#(mG)#(fC)#(mC)#(mC)#(mU)#(mU)#(fU) | (mG)#(mC)#(mA)(fU)(mC)(fA)(mU)(fC)(mA)(fA)(mU)(mU)(mU)(fC)#(mG)#(mA)-TegChol |
| In vivo Experiments | | |
| SOD1_123 Di-valent Original | V(mU)#(fC)#(mG)(fA)(fA)(fA)(mU)(fU)(mG)(fA)(mU)(fG)(mA)(fU)#(mG)#(fC)#(mC)#(mC)#(mU)#(fG)#(mU) | (mG)#(mC)#(mA)(fU)(mC)(fA)(mU)(fC)(mA)(fA)(mU)(mU)(mU)(fC)#(mG)#(mA)-DIO |
| SOD1_123 Methyl Rich | V(mU)#(fC)#(mG)(mA)(mA)(fA)(mU)(mU)(mG)(mA)(mU)(mG)(mA)(fU)#(mG)#(fC)#(mC)#(mC)#(mU)#(fG)#(mU) | (mG)#(mC)#(mA)(mU)(mC)(fA)(fU)(fC)(mA)(fA)(mU)(mU)(mU)(mC)#(mG)#(mA)-DiO |
| SOD1_123 exNA | V(mU)#(fC)#(mG)(fA)(fA)(fA)(mU)(fU)(mG)(fA)(mU)(fG)(mA)(fU)#(mG)#(fC)#(mC)#(mC)#(mU)#(exmU)#(exfU) | (mG)#(mC)#(mA)(fU)(mC)(fA)(mU)(fC)(mA)(fA)(mU)(mU)(mU)(fC)#(mG)#(mA)-DIO |
| SOD1_123 exNA low PS | V(mU)#(fC)#(mG)(fA)(fA)(fA)(mU)(fU)(mG)(fA)(mU)(fG)(mA)(fU)(mG)(fC)(mC)(mC)(mU)#(exmU)#(exfU) | (mG)#(mC)#(mA)(fU)(mC)(fA)(mU)(fC)(mA)(fA)(mU)(mU)(mU)(fC)#(mG)#(mA)-Dio |
